# Cross-species molecular stratification enables T cell receptor diagnosis of Parkinson’s disease

**DOI:** 10.64898/2025.12.15.694504

**Authors:** Jiayu Chen, Xian Xia, Yuruo Zhang, Xiaoge Tong, Yue Liang, Xiangding Fan, Qinghao Meng, Yihong Wei, Gang Hu

**Author notes:** These authors contributed equally to this work (Jiayu Chen, Xian Xia). Correspondence to (X.X.), (G.H.).

## Abstract

**Background:** Developing peripheral blood-based diagnostic models for idiopathic Parkinson’s disease (iPD), particularly those leveraging the T-cell receptor (TCR) repertoire, has long been considered infeasible because patient-derived TCRs appear to lack convergent sequence motifs. We reasoned that this apparent absence of shared TCR features likely reflects both insufficient sample sizes and unaccounted immune heterogeneity within the iPD population.

**Methods:** We reconstructed TCR repertoires from the two largest iPD cohorts currently available, anchoring them with three mechanistically distinct mouse models to stratify patients into model-anchored informed subtypes. Within these data-driven subtypes, we trained subtype-specific, multimodal multi-instance classifiers.

**Results:** We demonstrated that iPD patients can be successfully stratified into model-anchored informed subtypes. Furthermore, the trained subtype-specific, multimodal multi-instance classifiers achieved an Area Under Curve (AUC) exceeding 0.8 for both subtypes.

**Conclusions:** Our findings underscore the critical role of disease stratification in enabling TCR repertoire–based modeling in neurodegenerative diseases.

## Background

Evidence implicating the involvement of adaptive immune system in Parkinson’s disease (PD) is growing, highlighted by the discovery of α-synuclein (α-syn) reactive T cells in patients and broad signatures of peripheral immune activation [1–3]. T-cell receptors (TCRs), which collectively form a repertoire that catalogs an individual’s history of antigen exposure, offer a principal window into disease-associated immune processes and potential early triggers [4]. The prospect of uncovering a disease-specific TCR signature is compelling, as it could reveal common molecular factors for idiopathic PD (iPD) and potentially other neurodegenerative diseases [5]. However, the search for a reproducible TCR biomarker in iPD has been challenging, with early findings often failing to be validated in subsequent studies across different cohorts and technologies [6,7].

The failure to identify a consistent TCR biomarker in iPD likely stems from two fundamental challenges. First, many studies have been limited by small sample sizes and have restricted findings to specific epitopes, which lack the statistical power to detect subtle but real immunological differences [1,3,8]. Second, and perhaps more importantly, iPD is not a monolithic entity but a syndrome exhibiting substantial molecular heterogeneity [9]. Pathogenic routes range from genetic predispositions, such as those related to α-synuclein, to environmental or toxic exposures that may mimic the effects of substances like 1-methyl-4-phenyl-1,2,3,6-tetrahydropyridine (MPTP) [10]. It is plausible that these distinct origins imprint divergent immune signatures. Consequently, pooling heterogeneous patient populations in analyses can dilute or even completely obscure subtype-specific signals, leading to inconclusive results.

To overcome the sample size limitation, we reconstructed TCRs from peripheral blood RNA sequencing (RNA-seq) from two major cohorts in PD. Then we developed a novel framework to stratify patients by integration human data with three etiologically distinct mouse models. This integrative approach revealed two robust patient subtypes, which enabled us to train a subtype-specific diagnostic model using TCR repertoires and RNA data.

## Methods

Detailed information on materials and methods is available as supplementary materials.

### Data collection

Human data were sourced from the Parkinson’s Progression Markers Initiative (PPMI) database and the Parkinson’s Disease Biomarkers Program (PDBP) cohort, available through the Accelerating Medicines Partnership Parkinson’s Disease (AMP PD^®^) program [11,12]. All data collection and processing were conducted in accordance with the protocols established by these initiatives. A multi-step filtering process was applied to define the final study cohorts. The inclusion and exclusion criteria were as follows:

Sex: Male participants constituted the primary analysis cohort. Female participants were analyzed separately as an independent supplementary cohort using the same inclusion criteria and analytical workflow to assess the reproducibility and generalizability of the findings across sexes.

Diagnosis: Only participants with a stable diagnosis of either Parkinson’s disease or healthy control across all available visits were included. Individuals with diagnostic changes or other neurological conditions were excluded.

Genetic Status: The iPD cohort consisted of individuals with no known PD-associated genetic mutations. Participants identified with SNCA^+^ mutations were segregated and analyzed as a distinct ‘SNCA cohort’.

Sample Quality: To ensure data reliability, only RNA samples with an RNA Integrity Number (RIN) ≥ 7 were retained for downstream analysis.

Age: The study cohort was limited to individuals ≥ 50 years of age to reflect the demographic characteristics of typical disease onset.

Medication use: The PPMI cohort comprised exclusively de novo, treatment-naïve PD patients. For the PDBP cohort, we included both unmedicated patients and those with a confirmed history of levodopa usage. This selection criterion was applied to encompass patients with established exposure to the primary pharmacological treatment for PD.

Following participant filtering, a specific sample selection strategy was employed for each cohort. In PPMI Cohort, for each eligible participant, we selected the sample collected at the time point closest to their initial PD diagnosis. This approach was chosen to capture the immunological state of the early disease stages, where T-cell reactivity is hypothesized to be maximal. In PDBP Cohort, a sample was randomly selected from the longitudinal cohort of each participant to ensure data point independence. This random selection was intended to balance the age distribution between the iPD and HC groups.

The analysis of external datasets was restricted to specific subsets to ensure cohort consistency. We included male subjects aged >50 years from the COPD cohort [13] and male subjects aged >40 years from the CMV cohort [14] to maintain an adequate sample size. Given the significantly higher sequencing depth of the CMV dataset, we restricted the analysis to the top 0.5% of clonotypes by frequency to achieve a repertoire size comparable to our primary cohort. For the GICA [15] and LUCA [16] datasets, downsampling was implemented by sorting TCRs based on frequency and extracting the top 563 clonotypes to serve as model input.

### Animal Models and Procedures

3-month-old male C57BL/6J mice were used for the MPTP and PFF models. All animals were housed in a specific pathogen-free (SPF) facility under a 12-hour light/dark cycle with ad libitum access to food and water. All animal experiments were performed in accordance with procedures approved by the Institutional Animal Care and Use Committee (IACUC) of Nanjing Medical University (ethic committee permission #21110060). Detailed Mice Model Establishment, behavioral protocols, immunohistochemistry procedures, and blood collection methods are provided in Supplementary Methods.

### Time-series robust DEG analysis

To identify genes exhibiting time-dependent expression patterns that differ between experimental groups (PD models vs. controls), we employed a consensus approach combining maSigPro and two distinct modeling strategies within the DESeq2 likelihood ratio test (LRT) framework. To ensure robustness and mitigate method-specific false positives, the final set of PD time-related genes was defined as the intersection of genes identified as significant by at least two of these analysis methods. The maSigPro package (v1.56.0) [17] was used to model temporal gene expression profiles. Gene counts were first normalized using the Trimmed Mean of M-values method [18] in edgeR [19]. MaSigPro fits a polynomial regression model for each gene under a negative binomial framework. A design matrix was defined with a second-degree polynomial (degree = 2). For DESeq2 (v1.24.0) [20] we applied two complementary approaches: a likelihood ratio test comparing a full model including an interaction term between model type and age against a reduced model lacking this term, and a spline-based approach modeling age using natural cubic splines to accommodate non-linear temporal patterns. All models were adjusted for technical covariates including library quality metrics, RNA Integrity Number (RIN), library batch, and the expression levels of Hba-a1 and mt-Co1 to correct for erythrocyte contamination and mitochondrial content, respectively. P values were adjusted for multiple testing using the Benjamini-Hochberg (BH) method. Detailed model specifications and significance thresholds are provided in Supplementary Methods.

### Pathway enrichment analysis

Gene Ontology (GO) [21] and Kyoto Encyclopedia of Genes and Genomes (KEGG) [22] enrichment analyses were performed on the time-series DEGs using the R package clusterProfiler [23]. We used all expressed genes as the background gene set. GO terms and KEGG pathways with a BH adjusted p-value < 0.05 were considered significantly enriched.

### Differential Expression Analysis with Immune Cell Fraction Correction

To enable a robust comparison between mouse model and human patient peripheral blood transcriptomes, we performed differential expression analysis while statistically correcting for variability in immune cell proportions. All differential expression analyses were conducted using the ‘limma-voom‘ method [24] in R. A linear model was then fitted for each gene using the following design formula:

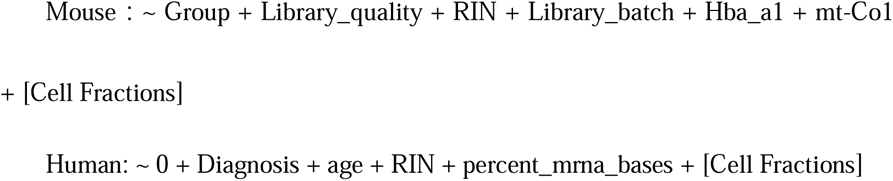

These cell fractions were computationally derived from the bulk RNA-seq data using CIBERSORTx [25]. To optimize the correction, different levels of cell-type granularity were evaluated (Fig. 3B).

### Cross-Species Transcriptomic Similarity Analysis

To systematically evaluate the translational relevance of our mouse models to human PD, we performed a multi-faceted similarity analysis between the mouse and human peripheral blood transcriptomes. We hypothesized that correcting for immune cell composition would enhance the concordance between species. Therefore, we compared results derived from three analysis schemes: 1) Naive: analysis without cell-type correction; 2) ICFR-main: correction for major immune lineages; 3) ICFR-fine: correction for fine-grained immune cell subtypes.

The similarity was assessed across three distinct dimensions. First, for Gene Expression Correlation of PD-Related Genes, we focused on a curated set of PD-relevant genes defined as all members of the "Parkinson’s disease" pathway in the KEGG database, and calculated the Spearman rank correlation coefficient between the log2 fold change of human PD compared to the control group and the corresponding log2FC in the three mouse models. Second, for Pathway Overlap Analysis using GSEA [26], we performed Gene Set Enrichment Analysis independently for the human and mouse datasets against three major pathway databases: Gene Ontology Biological Process (GO_BP), KEGG, and Reactome [27]. The degree of pathway overlap between species was quantified using the overlap index, calculated separately for the set of significantly upregulated pathways and the set of significantly downregulated pathways. Third, for Semantic Similarity, we computed the semantic similarity [28] between the enriched pathway sets from human and mouse for the GO_BP and KEGG pathways [29], using the GOSemSim [30] and simAnn [29] packages. To obtain a single metric of overall similarity, we applied the Best-Match Average (BMA) method [31]. Detailed parameters for GSEA thresholds and similarity calculations are provided in Supplementary Methods.

### MOUSEPAD integration

MOUSEPAD strategy is composed of the following four steps: (1) Learning latent manifold in mouse-human consensus space; (2) Identifying branches in latent space; (3) Annotating branches with etiologically informed mouse data; (4) Fitting trait dynamics, including TCR related traits, and cross-validating branches with existing PD subtypes. The analysis began from the ICFR-corrected mouse and human pathway level summary matrix.

For step (1), we first clustered samples with the Leiden [32] community detection algorithm and embedded the pathway matrix with three manifold learning algorithms: PHATE [33], UMAP [34], and t-SNE [35]. For step (2), to identify the axes of change, pseudotime analysis implemented in Palantir [36] was performed on the manifold. Palantir constructs a diffusion-based cell graph and defines a Markov chain over individual cells, computing pseudotime from local transition probabilities with entropy quantified as differentiation potential. To ensure the robustness of branch identification, we additionally applied VIA [37] in mouse-human consensus space. VIA is a graph-based algorithm that first clusters cells to build a kNN graph, then applies a lazy-teleporting random walk that preserves both local and global topology. Both algorithms employed the same starting point calculated as the sample with minimum medoid value across all samples.

For step (3), we evaluated mouse sample distribution on the manifold defined by PHATE and observed that time-series A30P and PFF samples tended to co-localize at one end of the PHATE embedding, while the MPTP samples were predominantly located at the opposite end. We then checked the chronological change of branch probability in the three mouse models and annotated the major two branches according to this result. For step (4), dynamics were calculated by regressing trait metrics to pseudotime in the two branches using a generalized additive model. We compared the MPTP branch with a previously phenotypically defined rapid-progress subtype [38] and the SNCA branch with SNCA mutation carriers in the PPMI dataset. Detailed algorithm parameters are provided in Supplementary Methods.

### Stability assessment for individual branch identity

To assess whether the branch variable represents an intrinsic attribute of each individual (i.e., if it is significantly associated with individual ID), we included more samples from the same participant in the original analysis, and a generalized linear mixed model (GLMM) was fitted. In this model, branch served as the binary response variable (SNCA or MPTP branch), and individual ID was included as a random intercept, as the followings:

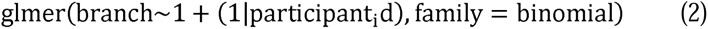

The estimated standard deviation of the individual random intercept was 0.89, with a 95% confidence interval of [0.685, 1.106]. As this confidence interval did not include zero, we concluded that the random effect of participant_id is statistically significant. This finding strongly supports the presence of significant intrinsic preference for branch status across individuals, rather than a purely random distribution.

We additionally checked the density distribution of random intercepts from the GLMM model, and the bimodal distribution indicated that the presence of latent heterogeneity across participants, which was consistent with branch identification.

### Independence of branch assignment from downstream classification

Branch assignments were derived exclusively from PD patients via unsupervised manifold learning and trajectory inference on KEGG pathway-level profiles; healthy controls were not involved at any stage of the MOUSEPAD pipeline and received no branch labels. The KEGG pathway-level input features represent each PD patient’s pathway activity as a deviation from the healthy baseline, such that the HC reference is effectively subtracted from the feature space prior to manifold learning. Branch labels therefore capture inter-PD heterogeneity in pathway perturbation patterns rather than PD-versus-HC discriminative information. Because healthy controls were entirely absent from the branch definition procedure, the downstream task of distinguishing PD from HC within each branch is orthogonal to the stratification step. Branch assignments were computed globally on the full PD cohort to ensure stable manifold estimation, and were subsequently treated as fixed, pre-computed attributes during all downstream cross-validation.

### Reconstructing the TCR repertoire from RNA-seq

The TCR repertoire was reconstructed from bulk RNA-seq data using TRUST4 [39] with default parameters. This tool assembles TCR CDR3 sequences by first identifying reads matching IMGT [40] gene segments (V, D, J, C) and then performing de novo assembly to build full-length receptor contigs. After annotation of the V and J genes, TRUST4 extracts the CDR3 sequence and its abundance for each identified clonotype.

### Repertoire diversity metrics

TCR repertoire statistics and diversity metrics were computed using Immunarch [41]. The total number of unique clonotypes was quantified, and CDR3 length distributions were obtained. Diversity estimators including Chao1 richness, Gini–Simpson index, Inverse Simpson index, D50 diversity, and Hill numbers were calculated. Clonal expansion was quantified through the proportion of the top 10 and top 20 most abundant clonotypes. V, J, and CDR3 amino acid usage frequencies were calculated directly from sample-level TCR annotations. Detailed formulas and function parameters are provided in Supplementary Methods.

### CDR3β sequence clustering and generation probability computation

To identify groups of TCRβ clonotypes with putative shared antigen specificity, we performed a large-scale clustering analysis. We aggregated the TCRβ repertoire data from our study with publicly available TCR datasets from peripheral blood of PD patients and controls, including GSE141578, GSE161192, β-chain TCR UMI counts [42] and TCR read counts [3]. This combined dataset was analyzed using the GLIPH2 algorithm [43,44] with default parameters, which utilize a reference dataset of mixed CD4+ and CD8+ TCRs. GLIPH2 clusters TCRs based on CDR3 sequence similarity (hamming distance of same-length CDR3) and motif enrichment of TCRβ clonotypes. We selected GLIPH2 similarity clusters that consisted of three or more unique CDR3 sequences and were present in three or more participants, with a Fisher_score ≤0.05, vb_score ≤0.05 and length_score ≤0.05. The disease association of these filtered clusters was assessed using a Fisher’s exact test (p ≤ 0.05) on a contingency table of their presence/absence in PD vs. HC subjects. Finally, to further characterize the TCRs within these clusters, we calculated their theoretical generation probability (P_gen_) for each unique CDR3β amino acid sequence using the Optimized Likelihood estimate of immunoGlobulin Amino-acid sequences (OLGA) algorithm [45]. The P_gen_ metric estimates the intrinsic likelihood of a sequence being generated via V(D)J recombination.

### Antigen prediction

To predict the antigenic targets of specific T cell receptor (TCR) clonotypes, TCR β-chain complementarity-determining region 3 (CDR3) amino acid sequences were compiled from two primary sources: (i) representative sequences from significant TCR similarity groups identified by the GLIPH2 algorithm, and (ii) top-ranking feature sequences derived from our MIL deep learning model. These compiled CDR3 sequences were submitted to the TCRmatch [46] online tool (http://tools.iedb.org/tcrmatch/) for alignment against experimentally validated TCR sequences curated in the Immune Epitope Database (IEDB) [47]. Following the tool’s documentation, a TCRmatch score greater than 0.97 was applied as the threshold to define a high-confidence match.

### TCR Sequence Embedding

To convert TCR sequences into numerical features suitable for machine learning, vector representations were generated using the TCR2vec deep learning framework [48]. Specifically, this framework, which is based on a 12-layer Transformer architecture, was employed to encode variable-length, full-length TCR sequences into uniform 120-dimensional embedding vectors.

### MIL deep learning model

To classify TCR repertoires based on disease status, a multiple instance learning (MIL) framework was employed [49]. In this paradigm, the entire TCR repertoire of a subject was treated as a bag, and each unique TCR sequence within it was considered an instance. A patient-level binary bag label was assigned to each repertoire, where bags from Parkinson’s disease (PD) patients were labeled as positive and those from healthy controls (HC) were labeled as negative.

To mitigate the influence of a large number of non-informative TCR instances within each repertoire, we adopted a Double-Tier Feature Distillation MIL (DTFD-MIL) architecture [50]. This model consists of a two-stage process involving pseudo-bag generation and hierarchical attention-based aggregation. First, for each bag, its instances were randomly partitioned into K smaller, non-overlapping pseudo-bags. Each pseudo-bag was assigned the original bag label. This step allows the model to learn from smaller, more manageable subsets of instances, effectively distilling key features. The core of our model is a gated attention-based MIL pooling mechanism [51], which was applied at both tiers. The DTFD-MIL framework operated via a two-stage, hierarchical attention process. In the first stage (Tier-1), the gated attention model was applied to each of the K pseudo-bags to generate a set of distilled feature vectors. In the second stage (Tier-2), these distilled vectors were treated as new instances and aggregated by a second attention model into a single, patient-level representation. This final vector was then passed to a fully connected layer with a sigmoid activation function for probability prediction. The entire model was trained end-to-end using a binary cross-entropy loss function. A systematic comparison of ten MIL methods spanning diverse methodological paradigms was performed (Supplementary Table 5). Detailed formulas and hyperparameters are provided in Supplementary Methods.

### RNA model and multimodal fusion

A Self-Normalizing Neural Network (SNN) [52] was implemented for the classification task, leveraging its ability to maintain normalized neuron activations throughout the network for stable training. The classifier architecture consisted of three fully connected hidden layers with Scaled Exponential Linear Unit (SELU) activation and alpha dropout. The final output layer consisted of a single neuron with a sigmoid activation function to produce a probability score for binary classification.

To integrate information from the TCR and RNA modalities and to model their synergistic interactions, a gating-based multimodal fusion strategy was used [53]. An attention-based gating mechanism was first applied to selectively emphasize informative unimodal features based on information from both modalities. The gated feature vectors were then fused using the Kronecker product to capture all bimodal interactions while preserving the original unimodal information. The resulting fusion tensor was flattened, L2-normalized, and propagated through two fully connected hidden layers for deep nonlinear processing of the fused features. The output of the final hidden layer was used for the downstream classification task. Detailed formulas and architecture specifications are provided in Supplementary Methods.

### Statistical analysis

All computational and statistical analyses were performed using the R (v4.2.3) [54]. For general comparisons of continuous variables, data are presented as mean ± standard error of the mean (SEM). Statistical significance was determined using two-tailed Student’s t-tests for two-group comparisons unless otherwise noted in the figure legends. For time-series RNA-seq data, differential expression was assessed using the Likelihood Ratio Test (LRT) within the DESeq2 framework and maSigPro, as described in the respective sections. The Benjamini-Hochberg (BH) method was applied to control the false discovery rate. A significance threshold of adjusted p-value < 0.05 was used as the default standard, unless otherwise specified in the relevant method sections or figure legends. Data visualization was generated using ggplot2 (v3.5.2) [55]. Loess regression lines were fitted with default parameters to visualize trends. Heatmaps and sequence logos were produced using pheatmap (v2.13.1) [56] and ggseqlogo (v0.1) [57], respectively.

## Results

### TCR motifs were enriched in PD patients but could not be directly used for diagnosis

To determine whether there existed global or local pattern difference between iPD patients and healthy controls (HC), we systematically collected and uniformly analyzed immune repertoire reconstructed from peripheral blood bulk RNA-seq (Fig. 1A, and Methods) collected by two large scale transcriptomic profiling studies, Parkinson’s Progression Markers Initiative (PPMI, iPD *n* = 229, HC *n* = 104) and Parkinson’s Disease Biomarkers Program (PDBP, iPD *n* = 212, HC *n* = 167) (Supplementary Table 1). We focused our analysis on male patients, guided by previous findings that α-syn-specific T cell responses were higher in males [6]. Then, to satisfy the independent and identically distributed assumption for statistical analysis, we considered each patient only once and random selection was restricted to early patients as previous study [6] found that α-syn-specific T cell response was negatively related to daily levodopa equivalent dose (Methods and Supplementary Fig. 1A).

**Figure 1.**
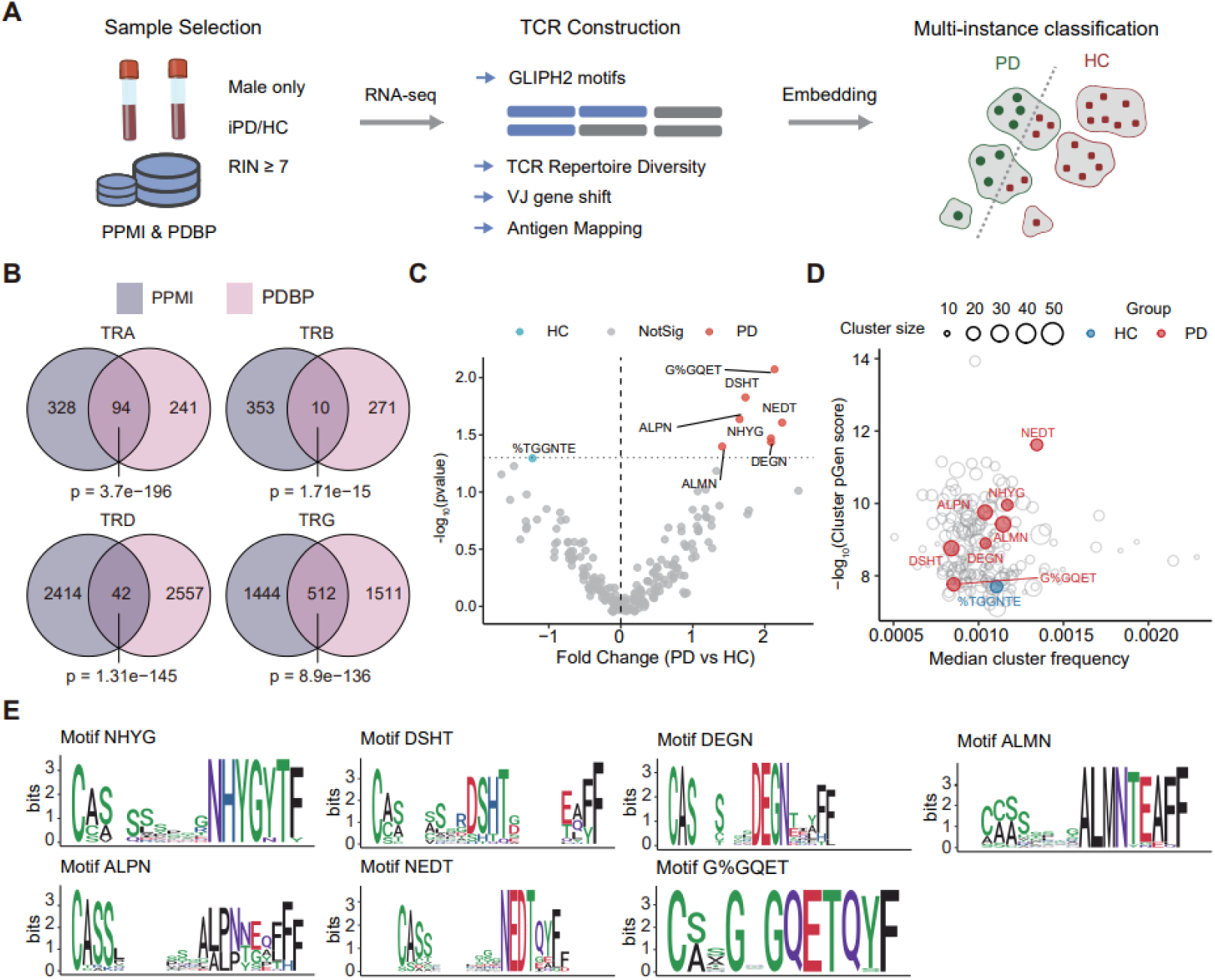
Characterization and identification of PD-associated T-cell receptor (TCR) signatures in PPMI and PDBP cohort. (**A**) Analysis workflow overview. TCR repertoires were reconstructed from selected AMP-PD cohort samples using TRUST4 [39]. The resulting repertoires were analyzed for differential features, which were subsequently used to train a deep learning-based classification model. (**B**) Venn diagram illustrating the overlap of public (clonal frequency > 0.0001) CDR3β sequences between iPD patients from PPMI and PDBP cohorts. The statistical significance of the overlap was calculated using Fisher’s exact test. (**C**) Volcano plot displaying the enrichment of TCR specificity groups in PD patients versus HC. TCRs were clustered into similarity groups using GLIPH2, and groups passing filtering criteria (*n* = 206) are shown. Each point represents a TCR group, colored by its significant enrichment in either the PD (red) or HC (blue) cohort as determined by Fisher’s exact test. The ‘%’ symbol indicates a wildcard (any amino acid) position within a shared motif. (**D**) V(D)J recombination generation probabilities (P_gen_) of CDR3 sequences belonging to the 206 significantly enriched GLIPH2 motifs from (C). Distributions are shown separately for motifs enriched in PD (red) and HC (blue), with high-P_gen_ motifs representing “public” TCR features that are more likely to be shared across individuals due to convergent recombination bias. (**E**) Peptide sequence logos the top 7 TCR motifs with the most abundant shared PD-specific CDR3 sequences.

We first ensured that the total TCR counts and the number of unique clones were comparable between iPD and HC across the four TCR types (Supplementary Fig. 1B).

Repertoire diversity metrics including Chao1 estimators, Gini-Simpson indices and inverse Simpson indices were not significantly different between iPD and HC in both cohorts for *TRA*/*TRB*/*TRD*/*TRG* (Supplementary Table 2), although we found iPD patients had significantly lower Gini-Simpson index (p = 0.037) and inverse Simpson index (p = 0.04) in *TRD* (Supplementary Fig. 1C) when considering samples from both genders. As previous report on TCR repertoire in α-synuclein-specific T cells showed surprising diversity [3], the overall identical pattern of *TRA/B/G* in iPD versus healthy control was not surprising, while the divergence in *TRD* diversity among all samples was unexpected and suggests potential gender-related effects on *TRD* composition. Unfortunately, current immunoinformatics tool regarding the TCR chains besides TCRβ chain was still lacking, so we could not go further on the biological insights of this difference. After false discovery rate correction, no significant differential usages of V(D)J genes were detected in all TCR chains of iPD patients (Supplementary Table 3), but at the gene family level, a usage preference of the *TRBJ* gene family toward *TRBJ2* in iPD patients was found (p = 0.007, Supplementary Fig. 1D). In particular, iPD patients displayed a higher frequency of *TRBJ2-7* and a lower frequency of *TRBJ1-6* usage (Supplementary Table 3). Furthermore, the distribution of CDR3 lengths did not differ among all TCR chains after FDR correction. We did observe significant number of common CDR3s across all TCR types (Fig. 1B) which indicated that iPD patients did share local feature similarity. In summary, here we systematically and rigorously queried TCR repertoire level features from two large scale peripheral blood transcriptome datasets and found that the global pattern of iPD patients was not significantly differ from healthy samples, but the local similarity existed in disease samples.

To take a further look at the local features on TCR sequences, we leverage the Grouping of Lymphocyte Interactions by Paratope Hotspots version 2 (GLIPH2) [44] to detect the specific enriched clusters and motifs in iPD patients. We identified 206 significant motifs after applying standard filtering criteria (Supplementary Table 4). Stratification by clinical group showed that seven motifs were significantly enriched in PD patients, whereas one was enriched in HC participants (Fig. 1C). We also applied GLIPH2 analysis to the female subset within the AMP-PD cohorts. In female patients, 8 TCRβ motifs were significantly enriched in PD compared to HC (Supplementary Fig. 2A). Notably, these 8 motifs showed no overlap with the 7 male-enriched motifs. However, when examining all motifs by directional trend (PD-skewed or HC-skewed regardless of individual significance), 20 PD-skewed motifs and 8 HC-skewed motifs overlapped between sexes (Supplementary Fig. 2B). This pattern suggests the presence of convergent, shared TCR features between male and female patients. We then assessed their generation probability and revealed that distinct clusters of motifs possess varying degrees of recombination likelihood (Fig. 1D, Methods). By integrating the probability of enrichment in iPD patient and the probability of natural generation, we found motif “G%GQET” as the both public and selected motif across iPD patients (Fig. 1E), and part of this motif “GQET” was recently reported in an independent study in single-cell TCR-seq from PD peripheral blood [58]. Taken together, we scrutinized the local enrichment of TCR sequence among iPD patients against healthy controls and we proved that with improved sample size, enriched motifs could be identified in iPD condition.

With common motifs identified in iPD patients, we further reached out for diagnostic models to differentiate iPD patients from healthy donors based on immune repertoire. We have compared 10 methods based on machine learning especially deep learning (Supplementary Table 5) [50,51,59–66] and most methods were based on β chain only. These ten benchmarked methods comprised two operational categories: four TCR-specific models that directly received raw TCR sequences (including DeepTCR, DeepRC, Mal-ID and DeepLION2), and six general-purpose MIL architectures that received dense TCR2vec embeddings as input (including DTFD-MIL, TransMIL, ABMIL, ACMIL, BiFormer and DSMIL). The best classification Receiver Operating Characteristic - Area Under the Curve (ROC-AUC) among all tested models was still only 0.56 (Supplementary Fig. 1E). We tested whether this can be attributed to lower number of TRB reads in RNA-seq reconstructed immune repertoire by down-sampling TCR-seq from two cancer datasets with distinct clonal architectures: a lung cancer cohort (LUCA) [16] (original TCR clones= 11491, AUC= 0.9615) characterized by private, patient-specific clonotypes, and a gastrointestinal cancer cohort (GICA) [15] (original TCR clones = 10505, AUC= 0.7179) enriched for shared-antigen-driven clonotypes. Down-sampling both to the depth in iPD repertoire (mean TCR clones = 563, Methods) led to markedly different outcomes reflective of their clonal architectures: the private-clonotype-dominant LUCA dataset suffered a substantial AUC reduction to 0.7435 (Δ = −0.218), whereas the shared-clonotype-dominant GICA dataset was largely preserved at 0.679 (Δ = −0.039) (Supplementary Fig. 1F). Critically, both retained classification performance well above chance, ruling out the possibility that shallow sequencing alone accounts for the poor performance of a unified TCR classifier in iPD.

### Generation of pathogen-specific time-series mouse reference transcriptome landscape

As the specific pathogenesis of PD remains elusive, and iPD patients exhibit distinct disease progression subtypes [38,67], we speculated that this etiological and clinical heterogeneity might preclude the existence of common TCR clusters among all patients, making a uniform repertoire-based classification model infeasible. Since dissecting specific etiological factor is challenging in human cohort studies but more tractable in mouse models, we hypothesized that projecting human peripheral blood transcriptome to mouse landscape across multiple models could reveal the subtypes from etiological aspect and find the conserved trajectory in iPD between mouse and human, thereafter to define the trajectory-specific repertoire-level PD classifier (Fig. 2A). To this end, we utilized three mouse models: (1) a neurotoxin-based model (chronic MPTP administration), (2) a genetic-based model (α-synuclein A30P mutation) and (3) an α-synuclein preformed fibrils (PFF) intrastriatal injection mode. Peripheral blood transcriptomes were sampled at four time points per model to capture dynamic changes across disease courses. Those mouse models were selected to represent different axes of PD etiology and pathology as genetic versus neurotoxic (A30P/PFF versus MPTP), brain-first versus whole-body (PFF versus A30P/MPTP) (Fig. 2A). We reasoned that they would collectively provide a representative reference for deconvolving heterogeneity within the human iPD population.

**Figure 2.**
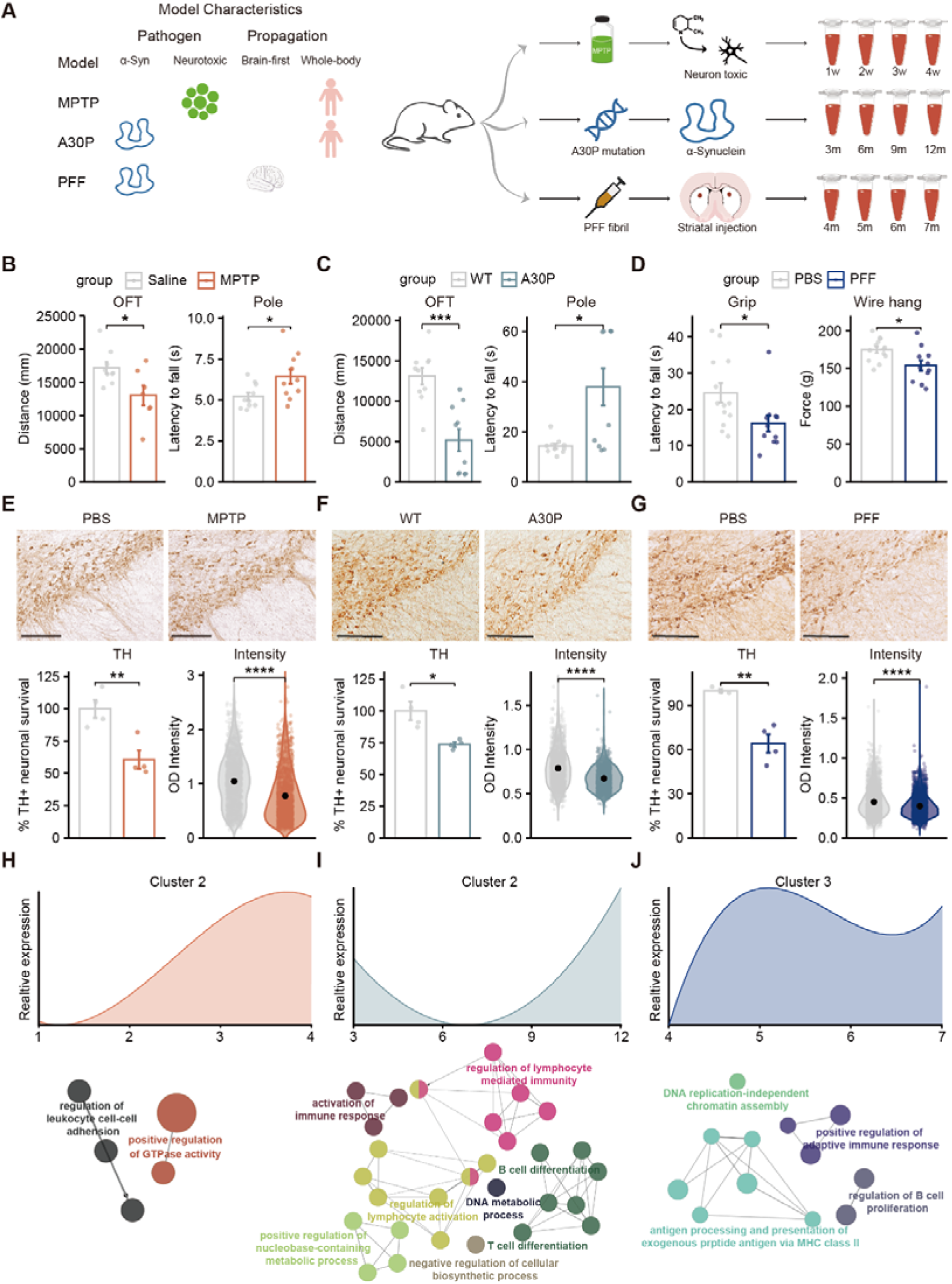
Pathological validation and temporal peripheral blood transcriptomic profiling of three distinct Parkinson’s disease mouse models. (**A**) Schematic overview of the experimental design. The left panel summarizes the key characteristics of the three models, highlighting their distinct pathogenic mechanisms and modes of pathology propagation. The right panel details the specific modeling procedures and longitudinal time points used for sampling peripheral blood, at each time point, 4–5 biologically independent samples were collected per group for each model and its corresponding control, followed by RNA sequencing. (**B-D**) Behavioral

**Figure 3.**
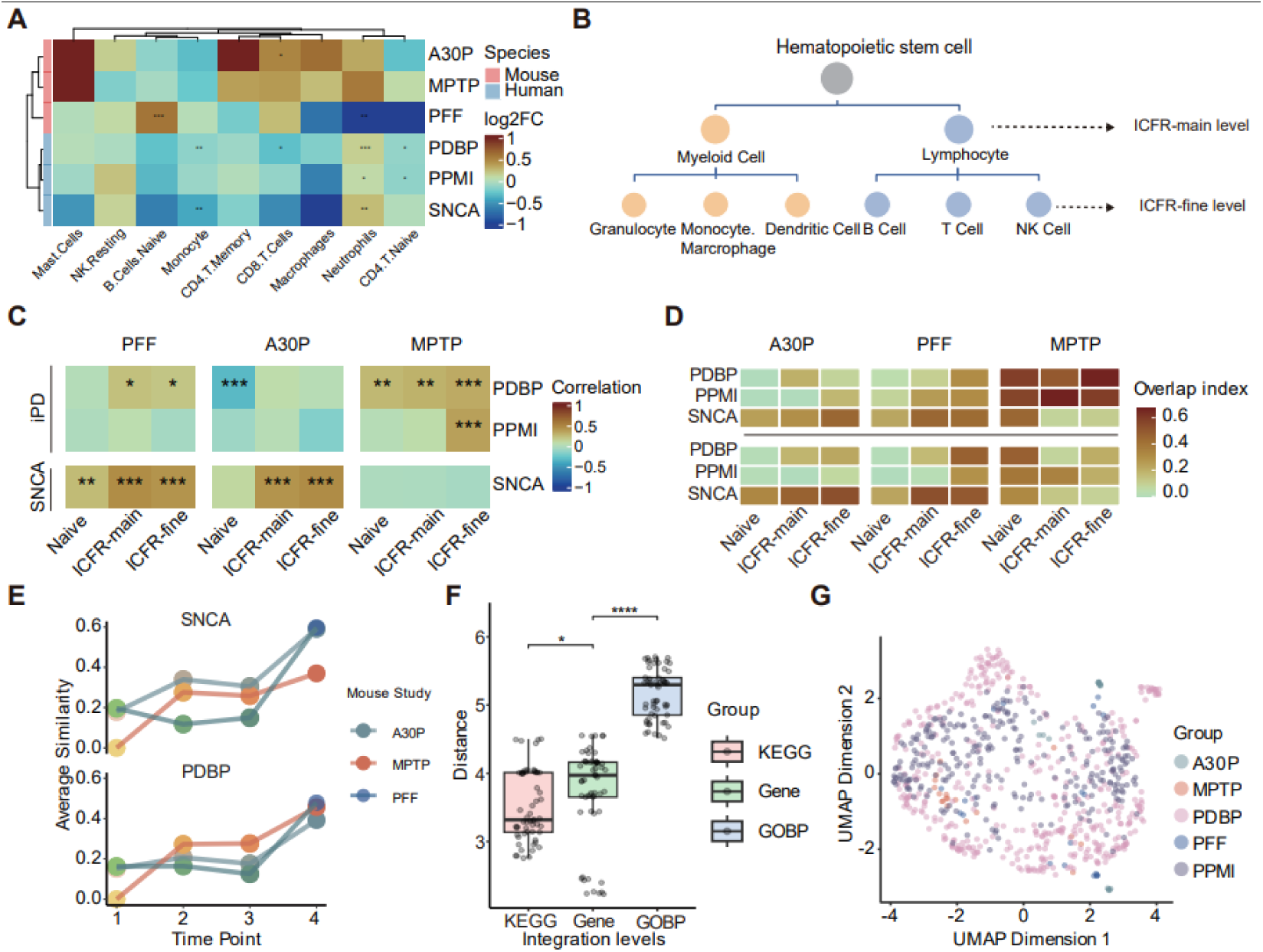
Cross-species integration of peripheral blood transcriptomes reveals conserved pathological pathways between PD mouse models and human patients. (**A**) Heatmap of relative changes in cell proportions in PD versus healthy controls (HC) across species. * indicates significant difference (t-test). (**B**) Schematic illustrating the cell-type hierarchy used for the immune cell fraction removal (ICFR) correction. The classification is based on the canonical hematopoietic differentiation tree for peripheral blood cells. (**C**) Spearman correlation analysis of logO(Fold Change) for genes within the KEGG “Parkinson’s disease” pathway. This analysis compares the expression changes in the three mouse models (MPTP, A30P, PFF) against those in three human patient cohorts (PPMI, PDBP, SNCA). (**D**) Pathway-level concordance between mouse models and human datasets. The overlap index quantifies the similarity of significantly upregulated pathways, shown for KEGG pathways (top) and Gene Ontology Biological Process (GO_BP) terms (bottom). Pathways were considered significantly upregulated if FDR < 0.25 and *p* < 0.05. (**E**) The plot shows that after ICFR-fine correction, the semantic similarity of shared KEGG pathways between mice and humans progressively increases over the course of the disease modeling, indicating a more biologically meaningful alignment as pathology develops. (**F**) Evaluation of integration effectiveness, measured by the average mouse-to-human distance in UMAP space at gene, GO_BP, and KEGG pathway levels. (**G**) UMAP projection of mouse and human samples based on their integrated KEGG pathway

To ensure mouse model quality, we assessed general altered motor functions in all three mouse PD models. Total distance from open field test showed impaired overall motor activity function in A30P (-60.4%) and MPTP (-32.0%) models (Fig. 2B and C), while no significant gross motor abnormalities was observed in PFF models (Supplementary Fig. 3A), consistent with its established progressive and prodromal behavioral phenotype [68]. Both A30P and MPTP models showed significant deterioration in the pole and rotarod tests (Fig. 2B and C and Supplementary Fig. 3B and C). While the latency to fall was decreased in PFF model, the abnormality was not significant (Supplementary Fig. 3A). So we further checked the muscular strength in PFF models with grip strength and wire hang test, and found they both yield significant decrease (-11.97% in grip strength, and -34.3% in wire hang, Fig. 2D), aligning with previous report [68]. Additionally, since constipation is the most common non-motor symptom that often precedes classical motor deficits [69], we measured fecal water content in PFF models and observed a significant reduction (Supplementary Fig. 3A).

We then quantified tyrosine hydroxylase (TH) positive cells in midbrain substantia nigra pars compacta (SNc) among all three models. In addition to severe loss of SNc TH^+^ neurons in all models (reduction of 26.4% in A30P, 39.4% in MPTP and 35.9% in PFF compared to age-matched controls), the expression level of TH was also decreased in the remaining neurons (Fig. 2E and G). Besides, we also observed accumulation of phosphorylated α-synuclein (pSyn, a marker of Lewy bodies or Lewy neurites), in the ventral area of SNc in PFF models (Supplementary Fig. 3D), consistent with previous report [68]. Taken together, we have established three PD models with distinct pathogenesis and the quality of each model was confirmed through comprehensive behavioral and pathological assessments.

To capture the trajectory of PD model through time, we collected peripheral blood across the time course among all three models followed by transcriptomic profiling (Fig. 2A). To investigate the temporal dynamics in PD mouse models relative to controls, we performed hierarchical clustering on the smoothed spline-based expression trajectories of robust differentially expressed genes (DEGs, Supplementary Fig. 4A and Methods), categorizing the genes into 4, 4, and 2 distinct clusters in the PFF, A30P, and MPTP models, respectively. Temporal clustering revealed distinct immunological signatures across the three models. In the α-synuclein-based A30P and PFF models, specific clusters showing sustained up-regulation (Cluster 2 in A30P; Cluster 3 in PFF) were enriched for B cell activation and antigen presentation, indicating a shared adaptive immune response (Fig. 2F and G). In contrast, the up-regulated cluster in the toxin-induced MPTP model was associated with innate inflammatory processes—including leukocyte adhesion, p38MAPK signaling, and cytokine production (IL-12, IL-10)—but showed no significant adaptive immune pathway enrichment (Fig. 2H). While other clusters across these models displayed variable patterns related to metabolism, translation, or homeostasis (Supplementary Fig. 4B), the divergence in major signaling pathways highlighted a clear distinction: α-synuclein manipulation elicited adaptive immunity, whereas MPTP pathology was dominated by innate inflammation. Consequently, aligning these distinct model signatures with human transcriptomic data could offer a strategy to recapitulate specific immune subtypes of the disease.

### Integration of representative mouse PD model and human profile

To molecularly stratify iPD patients, we implemented integration methods to maximize mouse-human similarity.in PD context. Because the proportions of peripheral blood cell types differ remarkably between mouse and human, global cross species comparisons remain challenging, despite extensive evidence of conserved features [70,71]. Moreover, shifts in cell type composition can generate large number of DEGs that obscure disease specific signals, whereas single cell analyses typically yield only modest DEG counts within individual cell populations [58,72]. Our data corroborated these established interspecies disparities [73], showing significant differences in overall cell distribution (Supplementary Fig. 5A). Furthermore, regarding disease-associated alterations, while neutrophil up-regulation was typically observed in human patients [72], we found a similar but non-significant trend in A30P and MPTP mice, whereas PFF mice exhibited downregulation (Fig. 3A), consistent with previous study[74]. These findings underscored the necessity of resolving cellular heterogeneity to ensure accurate cross-species comparison.

Then to enable integration of murine and human datasets, we applied immune cell fraction removal (ICFR, Methods) to eliminate the effect of cell proportions. We compared two cell removal strategies based on hematopoietic differentiation trajectories: the first strategy involved removal of myeloid and lymphoid lineages (named “ICFR-main”), while the second involved the exclusion of T cells, B cells, Natural killer (NK) cells, monocytes, and granulocytes (named “ICFR-fine”) (Fig. 3B). We evaluated the effectiveness of ICFR through gene- and pathway-level analysis. Although no concordance was observed among orthologous genes overall (Supplementary Fig. 5B) due to species-specific differences as indicated in previous study [75], we did observe positive correlation in PD-related genes (Fig. 3C, Methods). The correlation between A30P/PFF model and SNCA mutation carriers was even higher after ICFR-main and ICFR-fine correction (Fig. 3C and Supplementary Table 6), which was exceeding expectations because most PD-related genes were originally characterized in central nervous system studies and rarely interrogated in peripheral blood. Similarly, correlations also improved in comparisons between MPTP models versus iPD cohort after ICFR (Fig. 3C, Supplementary Table 6). Overall, ICFR-fine yielded higher overall correlations among two strategies.

At the pathway level, we performed gene set enrichment analysis (GSEA) [26] with adopted cutoffs (FDRO<O 0.25, PO<O0.05) to capture subtle enrichment signals. We found that the overlap index between mouse and human iPD data increased following ICFR-main and -fine in both up- and down-regulated Kyoto Encyclopedia of Genes and Genomes (KEGG) [22] pathways (Fig. 3D and Supplementary Fig. 5C), and the conclusion remain unchallenged in Gene Ontology Biological Process (GO_BP) [21] (Fig. 3D and Supplementary Fig. 5C) and Reactome database [27] (Supplementary Fig. 5D). Consistent with gene-level observations, the A30P/PFF model demonstrated a stronger resemblance to SNCA mutation carriers, while the MPTP model showed a closer profile to iPD patients (Fig. 3D). Since ICFR-fine outperformed ICFR-main, subsequent analyses primarily focused on ICFR-fine corrected data.

To investigate whether the similarity between mouse and human was time-course dependent, we quantified pathway level similarity between mouse and human cohorts by computing semantic similarity of pathways [29,30] and observed that similarity gradually increased with progression of mouse modeling in both KEGG and GO database (Fig. 3E and Supplementary Fig. 5E), suggesting that mouse peripheral blood transcriptomes reflect the pathological progression of PD.

To identify the most suitable level of data integration between mouse and human, we compared three levels: gene expression, GO terms and KEGG pathways. We performed Uniform Manifold Approximation and Projection (UMAP) dimension reduction at each level and used the average distance between mouse and human samples as an evaluation metric. The results indicated that integration at KEGG pathway level provided the most effective integration across species (Fig. 3F and G). In summary, these findings underscored the necessity of ICFR correction for comparative analyses of peripheral blood profile between mouse and human, and further revealed a time-dependent increase in similarity supporting the utility of time-series peripheral blood profiling for tracking disease progression.

### Immunology trajectories uncovered in PD via integrative transcriptomic manifold modeling

This comprehensive dataset was then subjected to a unified trajectory projection, forming a novel framework we termed Mouse-Anchored Pathway Analysis for Disease Trajectories (MOUSEPAD) (Fig. 4A). MOUSEPAD first integrated high-dimensional pathway data from both human and mouse samples. Subsequently, trajectory inference algorithms were applied to delineate disease progression paths. Finally, for each iPD patient, a precise position on the inferred trajectory was determined, including the probability of belonging to a specific branch and their pseudotime along different branches. ‘Trajectory’ here refers to a computationally inferred ordering in transcriptomic space and is not intended to imply a proven causal or temporal progression.

**Figure 4.**
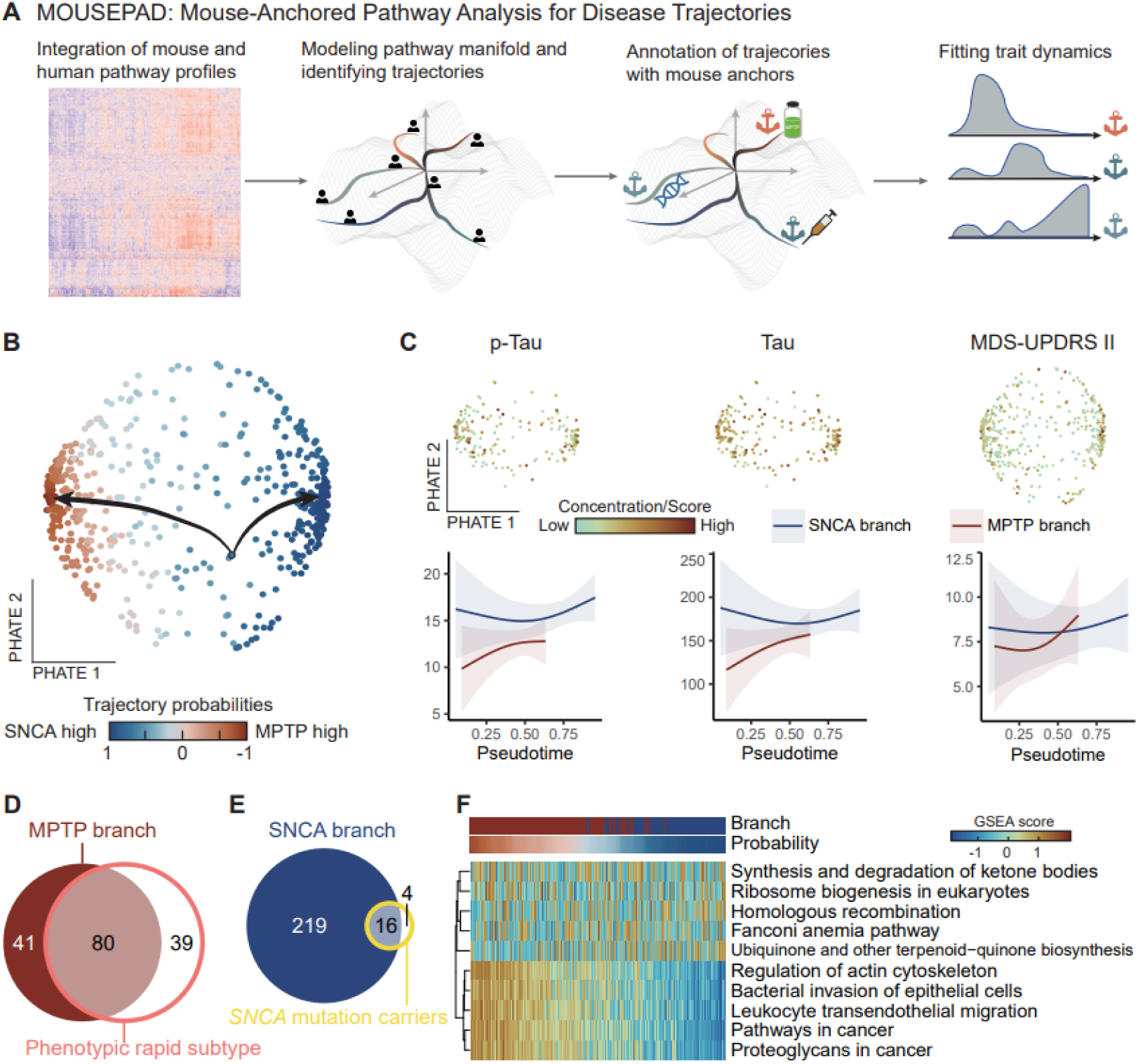
Modeling the mouse-human joint space manifold uncovered model-anchored informed trajectories of PD progression. (**A**) Schematics of the MOUSEPAD algorithm: (1) Learning latent manifold in mouse-human joint space; (2) Identifying branches in latent space; (3) Annotating branches with model-anchored informed mouse data; (4) Fitting trait dynamics, including TCR related traits, and cross-validating branches with existing PD subtypes. (**B**) The structure of the manifold capture by MOUSEPAD. A 2D PHATE embedding of each samples from mouse-human joint space. Dots are coloured by trajectory probabilities, and start point sample is circled. (**C**) Regression patterns of phenotypically change to pseudotime (left: p-Tau level in cerebrospinal fluid (CSF), pg/mL; middle: Tau level in CSF, pg/mL; right: Movement Disorder Society Unified Parkinson’s Disease

Upon applying the MOUSEPAD algorithm, we observed that iPD patients indeed segregated into two distinct branches (Fig. 4B). Crucially, time-series samples from the MPTP and SNCA-related (both A30P and PFF) mouse models of PD were predominantly distributed across these two branches, respectively (Supplementary Fig. 6A). Based on the distribution of the mouse anchor time-series samples, we provisionally designated these two paths as the SNCA branch and the MPTP branch.

These designations reflect molecular alignment with animal models that represent two orthogonal mechanistic axes: α-synuclein aggregation/propagation versus mitochondrial dysfunction/oxidative stress. They do not, however, imply a simplistic one-to-one mapping of multifactorial human iPD onto any single animal model.

To validate the robustness of this molecular subtyping, we first investigated potential differences in clinical data between those two branches. Analysis of cerebrospinal fluid (CSF) biomarkers revealed rapidly elevated levels of Aβ, total Tau, and phosphorylated Tau (p-Tau) in MPTP branch as compared to the stable trend on SNCA branch (Fig. 4C and Supplementary Fig. 6B). Clinical questionnaire assessments further demonstrated that patients on MPTP branch generally displayed more rapidly progressing motor and non-motor phenotypes (Fig. 4C and Supplementary Fig. 6B) than patients from SNCA branch. We also assessed the concordance between MOUSEPAD subtyping and previously established phenotype-based classifications from the PPMI and PDBP datasets. After implementing the methods of previous studies [38] to derive two phenotypic subtypes (Supplementary Fig. 6C), we performed enrichment analysis using our two MOUSEPAD branches. The result showed that MPTP branch was significantly overlap with phenotypic rapid subtype (Fig. 4D, Fisher’s exact test, odds ratio = 2.093, p = 0.008). Moreover, examination of SNCA mutation carriers within the PPMI dataset revealed their significant enrichment within the SNCA branch (Fig. 4E, Fisher’s exact test, odds ratio = 5.263, p = 0.002). Collectively, these results suggested that MOUSEPAD provided a reasonable molecular classification of iPD patients, and importantly, this peripheral blood transcriptome-based heterogeneity was strongly supported by converging phenotypic evidence.

We speculated that this molecular subtyping reflects fundamental differences in the underlying pathogenic trajectories of iPD patients. Therefore, we conducted an in-depth examination of the pathways making the largest contributions to these distinct trajectories (Fig. 4F). Up-regulation of ketone bodies, ribosome biogenesis and ubiquinone biosynthesis pathways suggested energy supply alteration on SNCA branch, while homologous recombination and fanconi anemia pathway indicated involvement of DNA repair. Up-regulation pathways in MPTP branch were more related to cell migration (pathways including actin cytoskeleton and leukocyte transendothelial migration) and infection related pathway, indicating the involvement of innate immunity. Additionally, we analyzed the distribution of median feature values from the raw projection matrix along the trajectory, finding a clear demarcation between the two branches (Supplementary Fig. 6D), suggesting an overall down-regulation of pathway median values in the SNCA branch and an inverse pattern in the MPTP branch.

To further ensure the stability of the MOUSEPAD classification, we employed alternative algorithms for trajectory projection, consistently observing that pseudotime inferences and probability estimation of two branches across different algorithms exhibited high correlation (Supplementary Fig. 6E and F). Given that both PPMI and PDBP datasets include longitudinal samples with multiple blood draws, we also investigated whether trajectory assignments were stable across different time points for the same sample (Supplementary Fig. 6G). The generalized linear mixed model confirmed that the branch classification was intrinsic for a cohort rather than random (95% confidence interval of [0.685, 1.106], Supplementary Fig. 6H), demonstrating the high stability of MOUSEPAD classification over multiple visits. To exclude the possibility that branch assignments were driven by cohort-specific or medication-related confounds, we examined the distribution of PPMI and PDBP patients across the two branches and found no significant enrichment by cohort origin (Supplementary Table 7, χ² test, p = 0.375) or by medication status within the PDBP cohort (Supplementary Table 8, χ² test, p = 0.817). Branch assignment is therefore independent of both cohort origin and treatment status. In conclusion, we introduced MOUSEPAD as a novel molecular subtyping system for iPD patients, highlighting its relevance to phenotypic subtypes and its potential molecular underpinnings. We proposed that this classification, anchored by distinct mouse PD models, could offer a more molecular informed understanding of peripheral immune signatures, hence facilitate the TCR diagnostic model building.

We also projected the female samples onto the MOUSEPAD framework derived from male mouse models. This branch assignment should be interpreted conservatively, given that our mouse anchors were generated exclusively from male animals and prior evidence indicates substantial transcriptomic heterogeneity between male and female PD patients [76]. Despite this limitation, female patients assigned to the fast-progression branch consistently exhibited more rapid worsening of both motor and non-motor phenotypes (Supplementary Fig. 7A). In addition, CSF biomarker trajectories (Aβ, total Tau, p-Tau) (Supplementary Fig. 7B) across the fast and slow branches showed directional consistency with those observed in the male cohort. These findings suggest that progression-based stratification is broadly reproducible across sexes at the phenotypic level, although the male-derived anchor model may not fully capture sex-specific molecular features of disease progression.

### Refined TCR-based deep learning diagnosis for PD

Finally, we examined whether the MOUSEPAD stratification of iPD patients indeed encompassed T cell–related immune features. First, leveraging motifs previously identified with GLIPH2, we compared motif loading across the two branches and observed higher disease-associated motif loading in PD patients within the SNCA branch than the MPTP branch (Fig. 5A). We further assessed differences in T cell subset composition and found significantly higher proportions of CD4 memory and CD4 naïve T cells in the SNCA branch relative to the MPTP branch (Fig. 5B), indicating branch-specific T cell differences.

**Figure 5.**
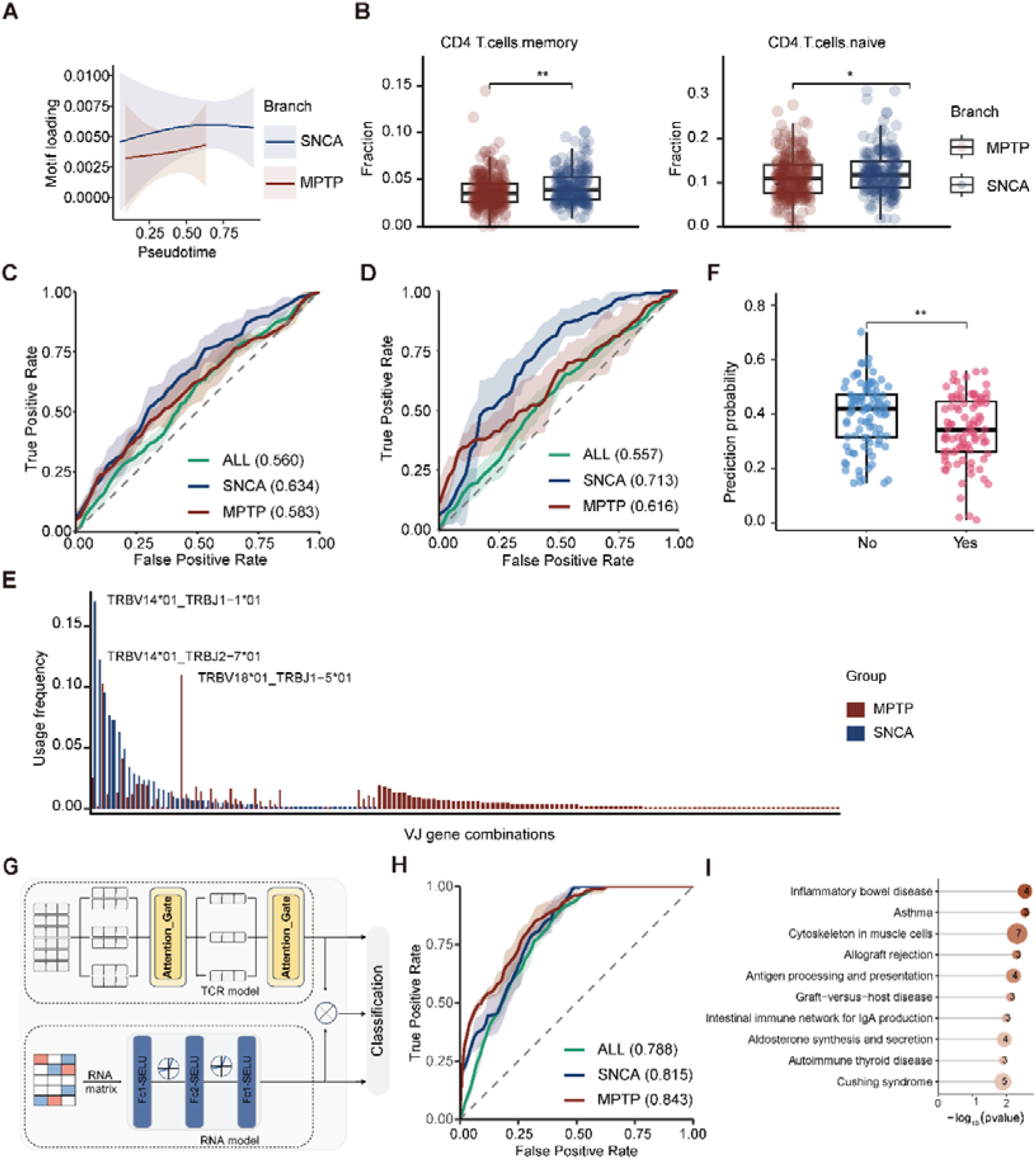
Classification and feature interpretation of the SNCA and MPTP branches of Parkinson’s disease. (**A**) Differential loading of PD-associated TCR motifs between the two PD branches. The plot shows that the aggregate loading scores of pre-identified PD-associated motifs are consistently higher in the SNCA branch compared to the MPTP branch. (**B**) Comparison of estimated T-cell subset proportions between the SNCA and MPTP branches. Bar plots display the relative proportions of CD4+ memory T cells (left) and CD4+ naïve T cells (right) for each branch. (**C**) Performance of the TCR-based classifier trained on the original AMP-PD dataset. ROC curves and corresponding AUC values are shown for the classification of the SNCA branch, the MPTP branch, and for a general PD vs. control classification (Overall). (**D**) Improved classifier performance after inclusion of diverse negative controls. ROC curves and AUC values are shown for the same classification tasks as in C), but for a model trained with additional COPD and CMV patient samples as negative controls. (**E**) V-J gene

We then developed a deep-learning classifier within a multiple instance learning (MIL) framework [50], which has been widely applied in medical imaging classification (Supplementary Fig. 8A). In five-fold cross-validation, the MIL model trained on TCR repertoire demonstrated higher performance for patients in the SNCA branch than for those in the MPTP branch (Fig. 5C): the AUC for the SNCA branch increased to 0.63, whereas the AUC for the MPTP branch remained at 0.58. We also applied the same DTFD-MIL classifier to the female TCR repertoire. The overall female model achieved an AUC of 0.59, modestly higher than the unstratified male model (AUC = 0.56). However, branch-specific performance differed substantially from the pattern observed in males. In females, classification performance was higher in the fast-progression branch (AUC = 0.642 ± 0.043) than in the slow-progression branch (AUC = 0.560 ± 0.015) (Supplementary Table 9). By contrast, in males, the slow-progressing SNCA branch outperformed the faster-progressing MPTP branch (AUC = 0.63 versus 0.58). This reversal indicates that the relationship between progression rate and TCR-based classification differed by sex. In females, the fast-progression branch showed higher classification performance than the slow-progression branch, whereas the opposite pattern was observed in males. Thus, progression-based stratification captured branch-specific signals in both sexes, but the branch hierarchy was not conserved across male and female cohorts. These findings suggest that branch assignments derived from male mouse anchors may not map directly onto equivalent biological states in female patients. Given the imbalance of healthy controls (*n* = 271) and PD patients (*n* = 441) in PPMI and PDBP cohort (Supplementary Fig. 1A), which contrasts with real-world distributions, we added age-matched male controls. Because many individuals in routine clinical settings present with non-neurological conditions and the PPMI recruitment did not exclude such comorbidities, we also incorporated other disease samples as controls; specifically, we introduced chronic obstructive pulmonary disease (COPD) samples (*n* = 202) [13] as additional negative controls. Importantly, these samples were used only in the training set and were not included in the test data. Adding the COPD dataset as negative control did improve the model’s performance, increasing the AUCs for the SNCA and MPTP branches to 0.675 and 0.619, respectively (Supplementary Fig. 8B). This effect was confirmed to be robust, as a similar performance gain was observed when using a separate cytomegalovirus (CMV) cohort as negative control [14] (Supplementary Fig. 8C). Ultimately, a combined model trained with both COPD and CMV controls achieved the highest performance, with the SNCA and MPTP branch AUCs reaching 0.713 and 0.616, respectively (Fig. 5D). We found that this sample augmentation strategy substantially improved discrimination in the SNCA branch, whereas only a limited improvement was observed in the MPTP branch.

To interpret the model’s predictions at a sample-specific level, we analyzed the attention scores generated by the model’s intrinsic attention mechanism. Based on the magnitude of these attention weights, we extracted the top three most impactful TCR features for each sample. Representative examples of these sample-specific predictive features identified via the attention mechanism are presented in Supplementary Table 10. To further characterize the biological properties of these prioritized features, and considering that TCRs recognizing the same antigen tend to exhibit biases in V-J gene usage [77], we first assessed V-J gene-usage differences (Fig. 5E). In the SNCA branch, TRBV14*01_TRBJ1-1*01 and TRBV14*01_TRBJ2-7*01 were the most frequently used V-J combinations, whereas TRBV18*01_TRBJ1-5*01 predominated in the MPTP branch. We also observed relatively concentrated V-J usage in the SNCA branch, in contrast to a broad, low-level distribution in the MPTP branch (Fig. 5E). Antigen prediction further indicated that 85.97% of PD-patient TCRs in the SNCA branch and 89.731% in MPTP branch were predicted to bind unknown antigens (Supplementary Fig. 8D and E).

Prior studies have reported associations between patient medication, age, and peripheral T-cell reactivity [6]. From the perspective of our model, we observed that the use of PD medications significantly reduced the predicted PD probability in the SNCA branch (Fig. 5F), with no detectable effect in the MPTP branch (Supplementary Fig. 8F), supporting the notion that therapy modulates TCR signals in PD. We did not detect a correlation between age and predicted probability (Supplementary Fig. 8G).

Finally, to better demonstrate the disease-discriminative potential of peripheral blood transcriptome in PD, we integrated TCR repertoire with expression data and constructed a multimodal model (Fig. 5G). The performance of RNA alone is provided in Supplementary Table 11. Upon integrating TCR and RNA modalities, the baseline AUC reached 0.788; after branch-specific modeling, the SNCA branch reached 0.81. Notably, although the MPTP branch underperformed on TCR features, it was better discriminated by RNA, yielding a final AUC of 0.84 (Fig. 5H). To interpret RNA feature contributions, we selected the top 200 protein-coding genes by feature importance and performed KEGG pathway enrichment. The SNCA branch features were significantly enriched in pathways of inflammation, immune disease, and antigen processing and presentation (Fig. 5I). This implicated a specific, systemic immune response as the dominant process in the SNCA branch. The involvement of antigen presentation pathways in peripheral blood suggested an active adaptive immune response, potentially targeting pathological α-synuclein or PINK1 protein and reflecting the systemic immunological nature of this synucleinopathy model. Conversely, the MPTP-branch features were enriched for pathways governing cell proliferation, migration, and metabolism (Supplementary Fig. 8H). This profile aligned with a systemic reaction to acute toxic insult. The metabolic pathway enrichment likely mirrored the mitochondrial dysfunction induced by MPTP in blood cells, while proliferation and migration pathways might represent a hematopoietic and immune cell response to systemic stress and tissue damage.

## Discussion

In this study, we conducted a comprehensive investigation into the peripheral immune landscape of Parkinson’s disease, leading to a novel patient stratification strategy and a high-performance, immune repertoire-based diagnostic model. We demonstrated that despite the subtlety of TCR repertoire alterations in the overall iPD population, these signals become potent diagnostic markers when viewed through the lens of disease heterogeneity. By developing a novel cross-species integration framework, MOUSEPAD, which leverages time-series transcriptomes from mechanistically distinct mouse models, we successfully stratified iPD patients into two subtypes with distinct levels of adaptive immunity, independent of known antigens and providing an unbiased classification. This stratification was the key that unlocked the development of a subtype-specific, deep learning-based classifier which, using only peripheral blood RNA-seq data, could accurately discriminate PD patients from controls.

In our initial analysis of TCR repertoires, reconstructed from bulk RNA-seq of two large cohorts (PPMI and PDBP), we observed that global repertoire metrics such as diversity and V(D)J gene usage were largely indistinguishable between iPD patients and healthy controls, corroborating the surprising diversity previously noted in α-synuclein-specific T cells [3].However, by integrating multiple datasets [3,42,78] to enhance statistical power, we identified seven TCR CDR3 motifs significantly enriched in iPD patients, indicating the existence of a subtle but convergent, public T-cell response. The inability of smaller studies to detect such motifs [3] underscores the necessity of large-scale, integrated analyses. The identified motif “G%GQET” exhibits a predicted P_gen_ exceeding 10^−8^, characterizing it as a high-frequency ‘public’ clonotype widely shared across the population [45]. Despite this ubiquity, it is significantly enriched within the PD cohort. Furthermore, this finding is independent corroborated by a recent single-cell study [58], thereby reinforcing the evidence for its disease association. Nevertheless, GLIPH2 identifies statistically enriched CDR3 similarity groups but does not infer antigen specificity. Consequently, these motifs should be interpreted as evidence of repertoire convergence rather than direct evidence for a shared disease-specific antigen. Despite these convergent signals, they were not sufficiently informative to support a simple unified diagnostic model; a comprehensive evaluation of multiple existing TCR-related classification algorithms [50,61–63,65,79,80] yielded poor classification performance. We confirmed through downsampling experiments that this failure was not attributable to the limited TCR read depth from RNA-seq, but rather pointed toward a more fundamental biological challenge.

This led us to hypothesize that the modest performance of a unified classifier stemmed from the well-documented molecular and clinical heterogeneity of iPD [81,82]. While prevailing subtyping efforts based on clinical or transcriptomic data have provided valuable insights [83–85], they have largely overlooked the underlying heterogeneity of peripheral immune responses. To probe this immune dimension, we turned to mouse models that, while only partially recapitulating the human disease [86], offer the ability to isolate specific pathogenic pathways. Our time-series transcriptomic analysis of the neurotoxic (MPTP), genetic (A30P), and α-synuclein propagation (PFF) models revealed distinct temporal immune signatures: the α-synuclein-driven models were characterized by a progressive adaptive immune response, consistent with an antigen-driven process [87,88], whereas the MPTP model displayed a predominant innate inflammatory profile [89]. In this framework, pseudotime does not represent developmental time or chronological age but rather a computationally inferred ordering of patient samples along axes of molecular disease progression, anchored by the time-series mouse transcriptomic data and validated by clinical phenotype concordance.

A major hurdle in translating these insights to humans is the profound difference in peripheral blood cell composition between species. Traditional cross-species comparisons often fail to account for these differences, leading to signals dominated by cell proportion changes rather than conserved disease biology [71]. To overcome this, we developed a novel computational approach, Immune Cell Fraction Removal (ICFR), which computationally corrects for compositional biases. This method significantly improved cross-species correlations at the functional pathway level, thereby enabling the creation of our MOUSEPAD framework. By projecting human patient data onto a manifold anchored by the distinct mouse immune trajectories, MOUSEPAD successfully segregated iPD patients into two robust subtypes: an “SNCA branch” aligned with the α-synuclein models and an “MPTP branch” aligned with the neurotoxin model. Crucially, this molecular stratification demonstrated significant clinical validity: the MPTP branch was enriched for patients with faster disease progression, while patients with known SNCA mutations were significantly enriched in the SNCA branch, confirming that our approach captured molecular relevant biology. Although cross-species transcriptomic similarity is inherently associative, the convergence of independent validation layers—from computational cross-species alignment metrics to human genetic anchors and clinical trajectory concordance—supports the biological coherence of these model-anchored molecular signatures beyond technical artifact. Rather than forcing a deterministic classification, the MOUSEPAD framework utilizes these distinct experimental models as biological lenses to deconstruct human molecular heterogeneity. Consequently, the framework is intrinsically architected to incorporate additional disease models as transcriptomic data become available, allowing this translational stratification to continuously evolve alongside our mechanistic understanding of PD pathogenesis.

Armed with this mouse model-anchored informed stratification, we revisited the challenge of TCR-based diagnosis. As hypothesized, T-cell-related immune features were significantly more pronounced in the SNCA branch, including an enrichment of disease-associated motifs and a higher proportion of CD4^+^ T cells, a subset previously shown to be reactive to α-synuclein epitopes [1,90]. This biological distinction enabled the training of a subtype-specific deep learning model that dramatically improved diagnostic accuracy for patients on the SNCA branch. The ultimate power of our approach was realized in a multimodal model that integrated both TCR repertoire and gene expression data. This combined model achieved high diagnostic accuracy across both subtypes, demonstrating strong potential as a clinically viable tool. Notably, both data modalities are derivable from a single peripheral blood RNA-seq assay, offering a promising, minimally invasive avenue for early diagnosis that is independent of traditional imaging or clinical assessments [91,92]. We observed that PD medication use significantly reduced the predicted PD probability exclusively in the SNCA branch (Fig. 5F). This branch specificity argues against a purely temporal confound, as disease duration alone would be expected to affect both branches. Dopamine receptors are functionally expressed on T cells and regulate T cell activation and proliferation [93]; levodopa and dopamine agonists may therefore directly dampen the α-synuclein-driven adaptive immune signals that characterize the SNCA branch. The α-synuclein-directed T cell response peaks in early, treatment-naive disease [6], and prodromal cohort samples may carry even more robust immune signatures before treatment-related dampening occurs.

Recent studies have identified several promising plasma biomarkers for PD, including neurofilament light chain (NfL) [19], plasma proteomic signatures [20,21], and blood-based α-synuclein seed amplification assays (SAAs) [22,23]. While these approaches have shown encouraging diagnostic performance, their generalizability across disease subtypes remains under investigation. In contrast, MOUSEPAD captures adaptive immune repertoire alterations and incorporates molecular stratification prior to classification, providing complementary information that may enhance future multimodal diagnostic frameworks for PD.

Our exploratory analysis of the female cohort revealed both similarities and divergences relative to the male-derived findings. Although a subset of PD-skewed GLIPH2 motifs showed directional concordance between sexes, the branch-stratified MIL classification exhibited an opposite pattern in females, with the Fast branch outperforming the Slow branch, in contrast to the male cohort. These results suggest that while certain TCR features are partially conserved across sexes, the MOUSEPAD-derived branch architecture, optimized on male-dominant data, does not fully capture female immune heterogeneity, highlighting the need for sex-stratified modeling and validation in independent female cohorts.

Our study has several limitations. First, our TCR repertoires were reconstructed from bulk RNA-seq, which provides shallower coverage than dedicated TCR-seq [97]. While we showed this was sufficient for subtyped classification, deeper sequencing may reveal additional informative features, and our down-sampling controls relied on DNA-based TCR-seq cancer datasets whose clonal architectures differ from the PD repertoire. Second, our analysis did not include patient HLA genotype information. Because T-cell responses are HLA-restricted, genetic variation in HLA loci across the population introduces significant heterogeneity. This may have masked more subtle, HLA-allele-specific TCR response signatures that could only be identified by stratifying patients according to their genetic background. Incorporating HLA data in future analyses could therefore refine the identification of public TCRs and further enhance the precision of diagnostic models. Third, our primary analysis was mainly limited to male participants to avoid the confounding variable of sex-related immune differences. Studies have shown that T-cell reactivity to PD-associated antigens and cell differentiation in female patients differ from those in male patients [8,98,99]. In exploratory analyses, sex-related differences were observed, suggesting that further validation in female cohorts and sex-specific models is warranted. Fourth, although our three mouse models were essential for establishing pathogenic subtypes, they only partially recapitulate the complex mechanisms of human PD. Fifth, although we identified disease-associated TCRs and their probable branch-specific activities, the specific antigens they recognize remain unknown; TCRMatch-based antigen prediction assigned matches predominantly to viral epitopes, with >85% of TCRs unassigned, and definitive antigen identification will require orthogonal experimental validation such as tetramer staining or functional T-cell assays. Finally, although our cohort represents the largest TCR repertoire analysis in iPD to date, the sample size remains modest by deep learning standards. Our strict filtering was designed to prioritize signal quality over sample quantity; validation in larger, independent cohorts with diverse demographic compositions will be essential to establish the generalizability of these subtype-specific classifiers.

In conclusion, our study demonstrates that deconstructing the heterogeneity of Parkinson’s disease through a cross-species, systems-immunology approach is key to unlocking the diagnostic potential of peripheral immune profiles. The MOUSEPAD framework provides not only a robust tool for molecular subtyping but also a clear path toward developing precise, blood-based diagnostics for this complex neurodegenerative disorder. By synergizing biological insights from mouse models with large-scale human data, our work solidifies a foundation for accurate, immune repertoire based diagnostics that fully account for the inherent complexity of neurodegenerative diseases.

## Conclusions

Our findings underscore the critical role of disease stratification in enabling TCR repertoire–based modeling in neurodegenerative diseases. The inherent molecular and clinical heterogeneity of idiopathic Parkinson’s disease obscures generalized peripheral immune signals, rendering universal diagnostic approaches ineffective. By deconstructing this complexity into biologically distinct subtypes, we demonstrate that resolving underlying trajectory differences is a mandatory prerequisite for capturing meaningful adaptive immune responses.

## Abbreviations

iPD: Idiopathic Parkinson’s disease
TCR: T-Cell Receptor
AUC: Area Under Curve
PD: Parkinson’s disease α-syn α-synuclein
MPTP: 1-methyl-4-phenyl-1,2,3,6-tetrahydropyridine
RNA-seq: RNA sequencing
PPMI: Parkinson’s Progression Markers Initiative
PDBP: Parkinson’s Disease Biomarkers Program
AMP-PD: Accelerating Medicines Partnership Parkinson’s Disease
RIN: RNA Integrity Number
PFFs: Pre-formed fibrils
HC: Healthy controls
GLIPH2: Grouping of Lymphocyte Interactions by Paratope Hotspots version 2
LUCA: Lung cancer
GICA: Gastrointestinal cancer
TH: Tyrosine hydroxylase
SNc: Substantia nigra pars compacta
pSyn: Phosphorylated α-synuclein
DEGs: Differentially expressed genes
ICFR: Immune cell fraction removal
GSEA: Gene set enrichment analysis
KEGG: Kyoto Encyclopedia of Genes and Genomes
GO: Gene Ontology
UMAP: Uniform Manifold Approximation and Projection
MOUSEPAD: Mouse-Anchored Pathway Analysis for Disease Trajectories
CSF: Cerebrospinal fluid
p-Tau: Phosphorylated Tau
MIL: Multiple instance learning
COPD: Chronic obstructive pulmonary disease
CMV: Cytomegalovirus

## Data availability

Mouse peripheral blood RNA-seq data are available at the National Genomics Data Center. AMP PD data and quality control notebooks are access-controlled (https://amp-pd.org/) and require individual sign-up to access the data.

## Code availability

The complete code used for the analysis in this study will be made publicly available on GitHub upon publication.

## Acknowledgements

Data used in the preparation of this article were obtained from the Accelerating Medicine Partnership® (AMP®) Parkinson’s Disease (AMP PD) Knowledge Platform. For up-to-date information on the study, visit https://www.amp-pd.org/. The AMP® PD program is a public-private partnership managed by the Foundation for the National Institutes of Health and funded by the National Institute of Neurological Disorders and Stroke (NINDS) in partnership with the Aligning Science Across Parkinson’s (ASAP) initiative; Celgene Corporation, a subsidiary of Bristol-Myers Squibb Company; GlaxoSmithKline plc (GSK); The Michael J. Fox Foundation for Parkinson’s Research ; Pfizer Inc.; AbbVie Inc.; Sanofi US Services Inc.; and Verily Life Sciences. Clinical data and biosamples used in the preparation of this article were obtained from the Parkinson’s Progression Markers Initiative (PPMI), and the Parkinson’s Disease Biomarkers Program (PDBP). Data used in the preparation of this article was obtained on 2022-08-01from the Parkinson’s Progression Markers Initiative (PPMI) database (www.ppmi-info.org/access-data-specimens/download-data), RRID:SCR_006431. For up-to-date information on the study, visit www.ppmi-info.org. PPMI – a public-private partnership – is funded by the Michael J. Fox Foundation for Parkinson’s Research funding partners 4D Pharma, Abbvie, Acurex Therapeutics, Aligning Science Across Parkinson’s, Allergan, Amathus Therapeutics, Avid Radiopharmaceuticals, Bial Biotech, Biogen, BioLegend, Bristol-Myers Squibb, Calico, Celgene, Dacapo Brain Science, Denali, The Edmond J. Safra Foundaiton, GE Healthcare, Genentech, GlaxoSmithKline, Golub Capital, Handl Therapeutics, Insitro, Janssen Neuroscience, Lilly, Lundbeck, Merck, Meso Scale Discovery, Neurocrine Biosciences, Pfizer, Piramal, Prevail, Roche, Sanofi Genzyme, Servier, Takeda, Teva, UCB, Verily, and Voyager Therapeutics. The Parkinson’s Disease Biomarker Program (PDBP) consortium is supported by the National Institute of Neurological Disorders and Stroke (NINDS) at the National Institutes of Health. A full list of PDBP investigators can be found at https://pdbp.ninds.nih.gov/policy. The PDBP investigators have not participated in reviewing the data analysis or content of the manuscript. We thank Yao Wei for his valuable study design advice, and Yinquan Fang and Xin Ding for their guidance on experimental techniques.

## Author contributions

X.X. conceived and designed the study. X.X. and J.C. organized data from AMD-PD and developed mouse PD model. J.C. implemented TCR analysis, ICFR and MIL/DL algorithms. X.X. implemented MOUSEPAD framework. Y.Z., Y.L., R.T., Q.M., and

Y.W. assisted in mouse model development. X.T. assisted in IHC. X.F. implemented RNA-level PD prediction. X.X. and G.H. acquired funding. X.X. supervised the study. X.X. and J.C. wrote and revised the manuscript, which was reviewed, edited, and approved by all authors.

## Ethics declarations

### Ethics approval and consent to participate

Animal experiments were approved by the Institutional Animal Care and Use Committee (IACUC) of Nanjing Medical University (ethic committee permission #21110060).

### Consent for publication

Not applicable.

