## Supplementary Figures for "Cross-species molecular stratification enables T cell receptor diagnosis of Parkinson’s disease"

**The PDF file includes:**

Fig. S1 to S8.

Supplementary Tables 1 to 9 and 11.

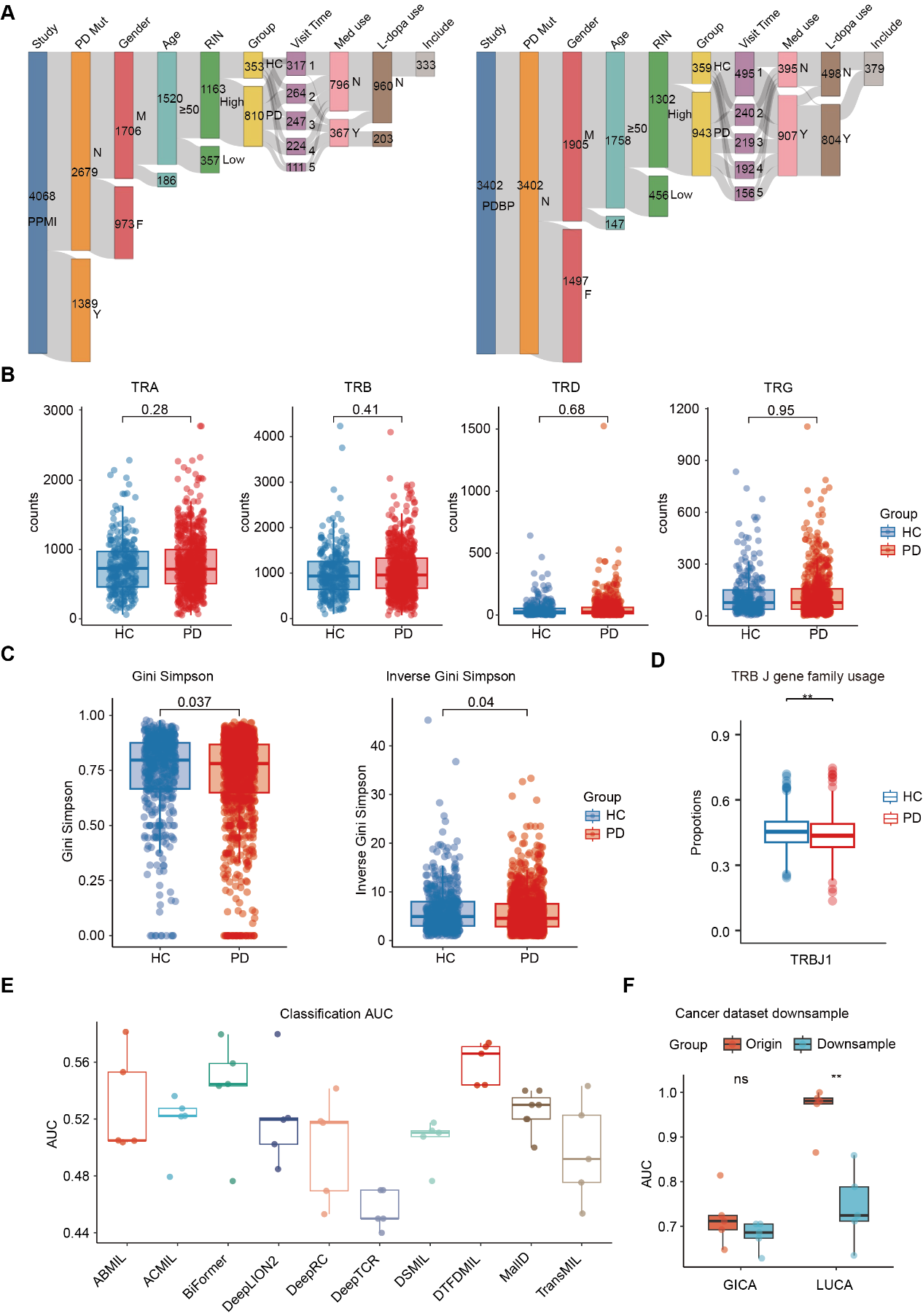

**Supplementary Figure 1. Overview of TCR repertoire characteristics and model performance in PD and HC cohorts.** (**A**) Sankey diagram illustrating the sample filtering process in the PPMI and PDBP cohorts. Numbers indicate sample counts at each step. Abbreviations: PD mut, PD mutation; Med use, PD medication use; L-dopa use, levodopa use; Y, yes; N, no; M, male; F, female. (**B**) TCR counts between PD and HC in TRA/B/D/G sequence. (**C**) Repertoire diversity (Gini-Simpson and Inverse Gini-Simpson indices) in the combined male and female PD and HC cohorts. Significance was determined by a student’s t test. Boxplots show the median and interquartile range. (**D**) Differential TRBJ gene usage in male PD patients versus HCs. (**E**) Boxplots of Area Under the Curve (AUC) scores for ten classification models evaluated using 5-fold cross-validation. (**F**) The AUC performance on the GICA and LUCA dataset and after downsampling the original TCR read counts to a similar number of TCRs as AMP-PD datasets. * *p* < 0.05, ** *p* < 0.01.

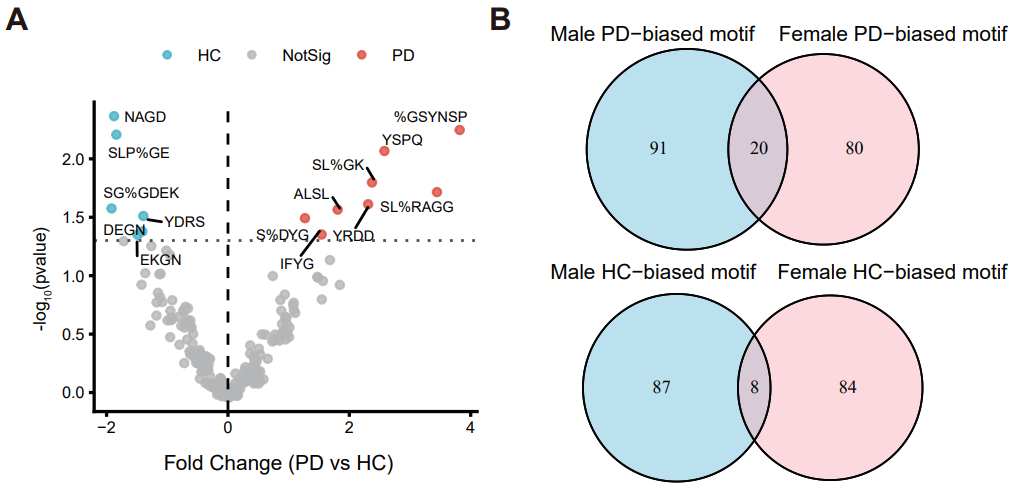

**Supplementary Figure 2. GLIPH2 analysis in female PD patients.** (**A**) GLIPH2 analysis of the female subset from the AMP-PD cohorts identified 8 TCRβ motifs significantly enriched in PD compared with healthy controls (HC). (**B**) Overlap analysis of TCRβ motifs stratified by directional enrichment (PD-skewed or HC-skewed regardless of statistical significance) between male and female cohorts. Twenty PD-skewed motifs and eight HC-skewed motifs were shared between sexes.

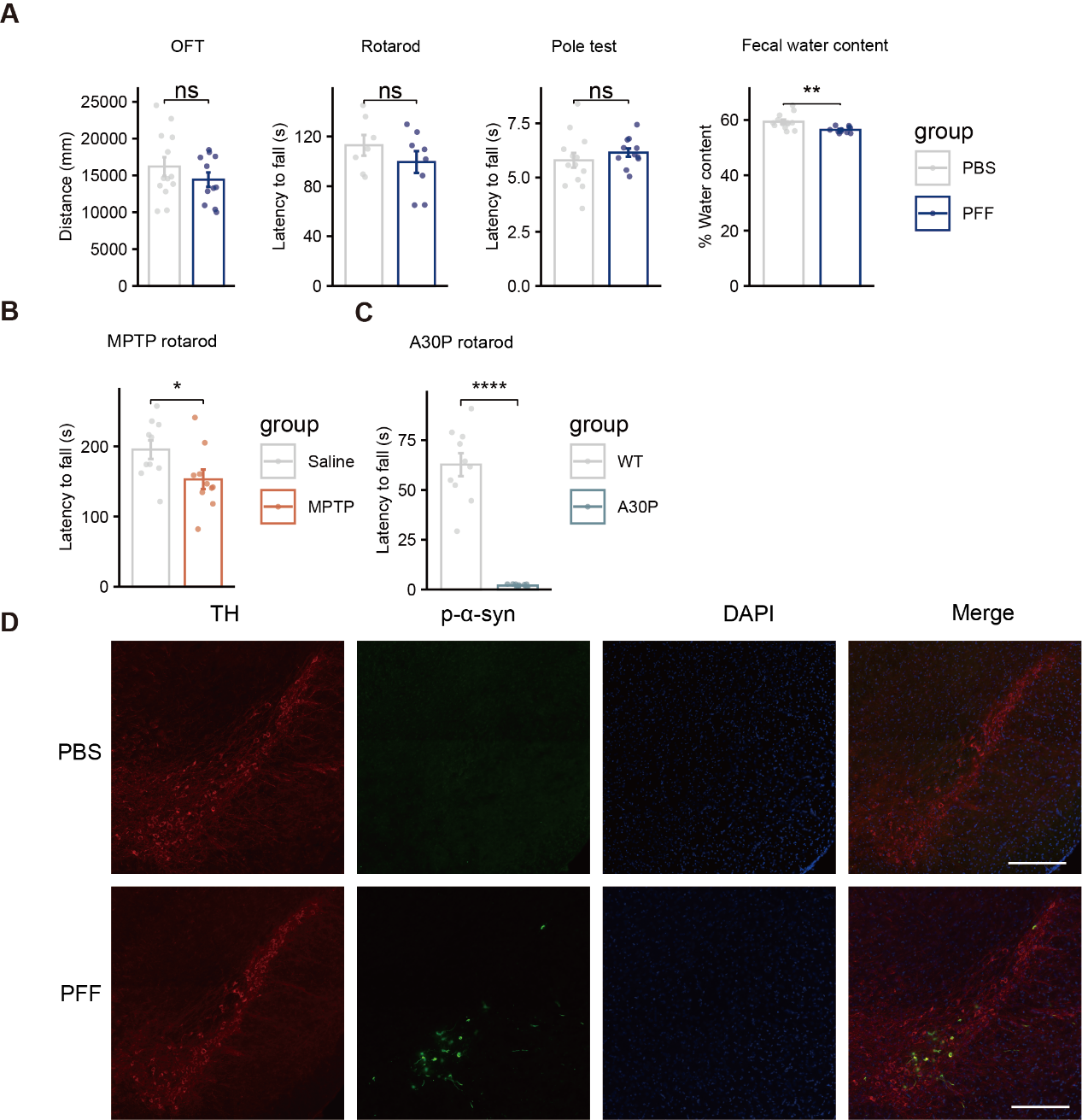

**Supplementary Figure 3. Behavioral assessments and dopaminergic neuron pathology in PD mouse models.** (**A**) Behavioral performance of the PFF model in the open field, rotarod, and pole tests, along with fecal water content analysis for non-motor symptoms. (**B-C**) Rotarod test performance for the (B) MPTP and (C) A30P mouse models. D) Representative fluorescence images of the SNc in PFF and PBS control mice. Data are presented as mean ± SEM; Student's t-test, **P* < 0.05, ***P* < 0.01, ****P* < 0.001.

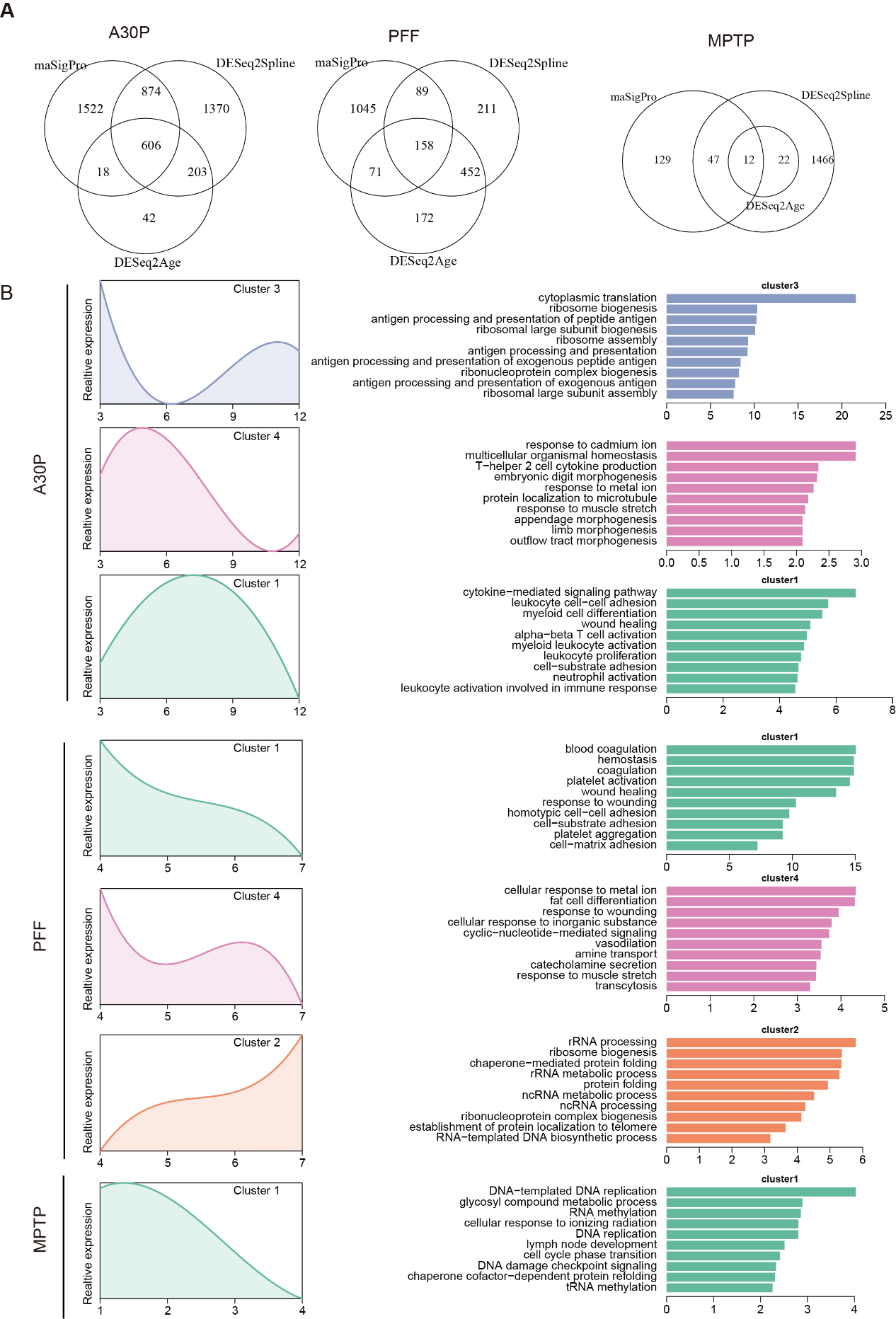

**Supplementary Figure 4. Shared temporal transcriptional programs across PD mouse models.** (**A**) Venn diagram showing the overlap of temporal differentially expressed genes (DEGs) identified in the A30P, PFF, and MPTP models. Temporal DEGs were defined by a consensus of three methods (maSigPro, DESeq Age, DESeq Spline). (**B**) Functional analysis of shared temporal expression clusters. Left panels display the expression dynamics of co-expressed gene clusters identified across all three models. Right panels show the top 10 significantly enriched GO Biological Process terms for each corresponding cluster, revealing the functional pathways associated with these shared temporal patterns.

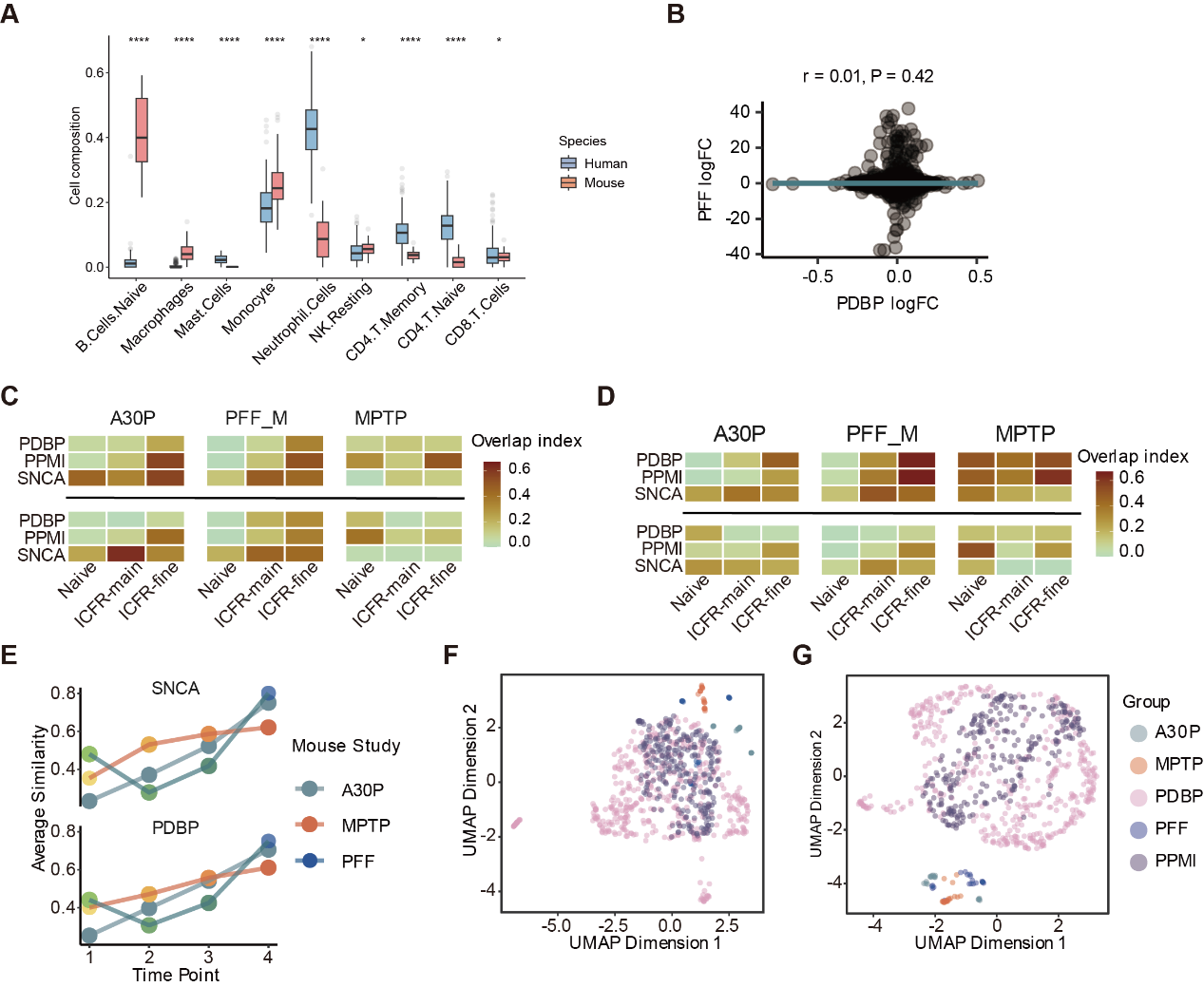

**Supplementary Figure 5. Cross-species concordance between mouse models and human PD blood transcriptomes.** (**A**) Differences in blood cell-type fractions between healthy human and mouse samples. (**B**) Spearman correlation of gene-level log₂FC values between the PFF mouse model and the human PDBP cohort, using all orthologs. (**C**) Overlap index of shared downregulated KEGG pathways (top) and GO BP terms (bottom) between mouse and human datasets (FDR < 0.25, p < 0.05). (**D**) Overlap index of shared upregulated (top) and downregulated (bottom) Reactome pathways between mouse and human datasets (FDR < 0.25, p < 0.05). (**E**) Increased semantic similarity of GO BP terms between mouse and human data after full ICFR-fine correction. (**F-G**) UMAP projections of mouse and human samples based on integrated profiles at the (F) gene level and (G) GO BP level.

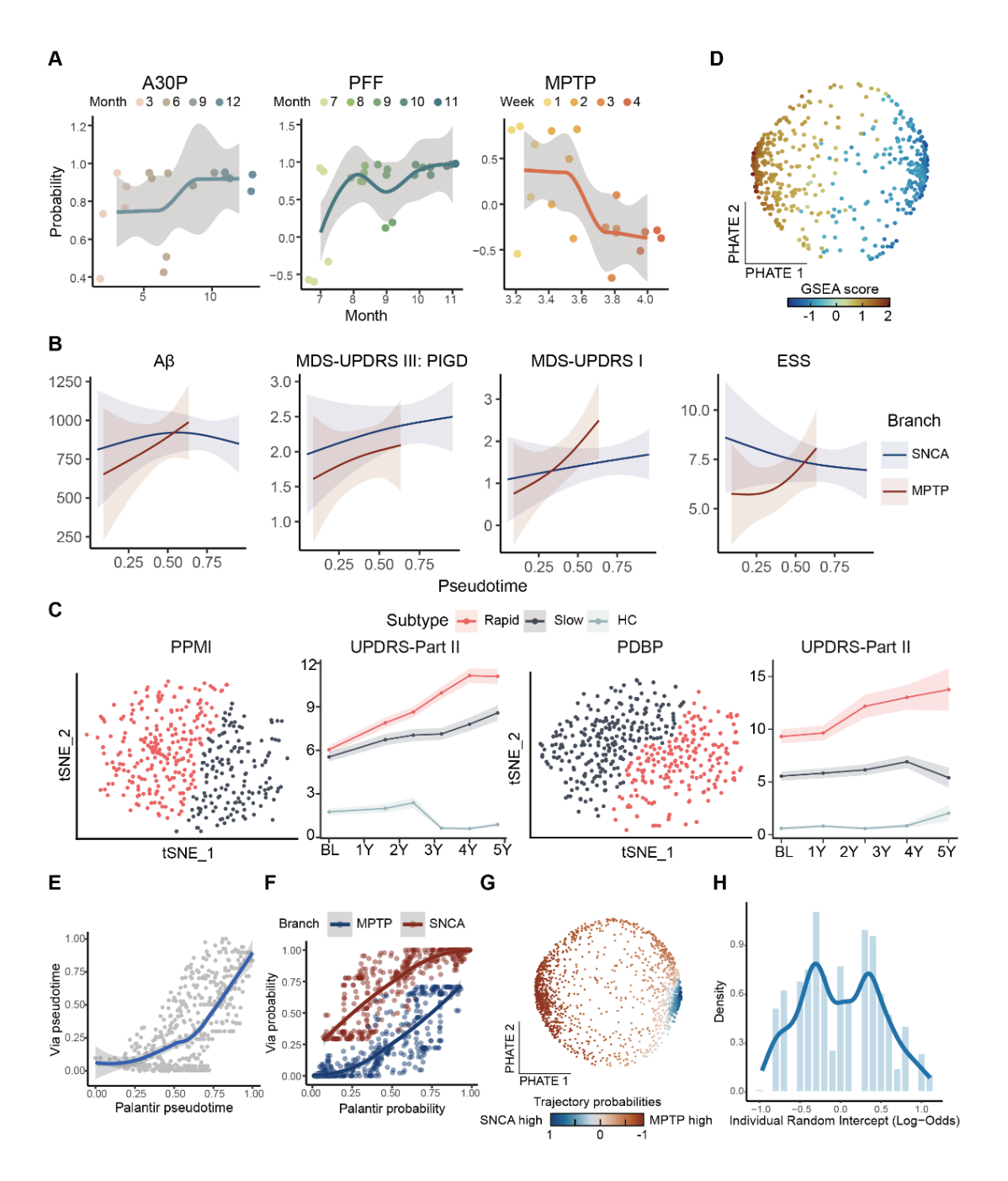

**Supplementary Figure 6. Mouse anchoring and robustness analysis of MOUSEPAD.** (**A**) The probability spanning of time series mouse corrected profile, showing the two SNCA-related mouse models are with positive relationship with branch probability while MPTP model is on the opposite direction. The regression was calculated with Loess regression and probability was derived from Palantir algorithm. (**B**) Additional regression patterns of phenotypically change to pseudotime (from left to right: Aβ level in CSF, pg/mL; MDS-UPDRS part III assessing postural instability and gait disorder (PIGD) score for additional motor phenotype; MDS-UPDRS part I score and Epworth Sleepiness Scale (ESS) total score assessing non-motor phenotypes) along the two trajectories (Methods). The error bands show the 95% confidence intervals. (**C**) Subtypes defined by PACE algorithm. PPMI (left) and PDBP (right) were processed separately because the input data could not be unified. Both subtype definition (left on each dataset) and corresponding phenotype progression (right on each dataset) were shown. (**D**) Distribution of median GSEA score on PHATE manifold. (**E-F**) Association between Palantir and VIA on pseudotime estimation (E, Pearson’s correlation = 0.67) and branch probability (F, Pearson’s correlation = 0.85). (**G**) Samples from the same participants but on different time points were included in this latent manifold learning. Branch probability were visualized to show two branches still holds for expanded sample size. (**H**) Density distribution of random intercept derived from generalized linear mixed model (Methods).

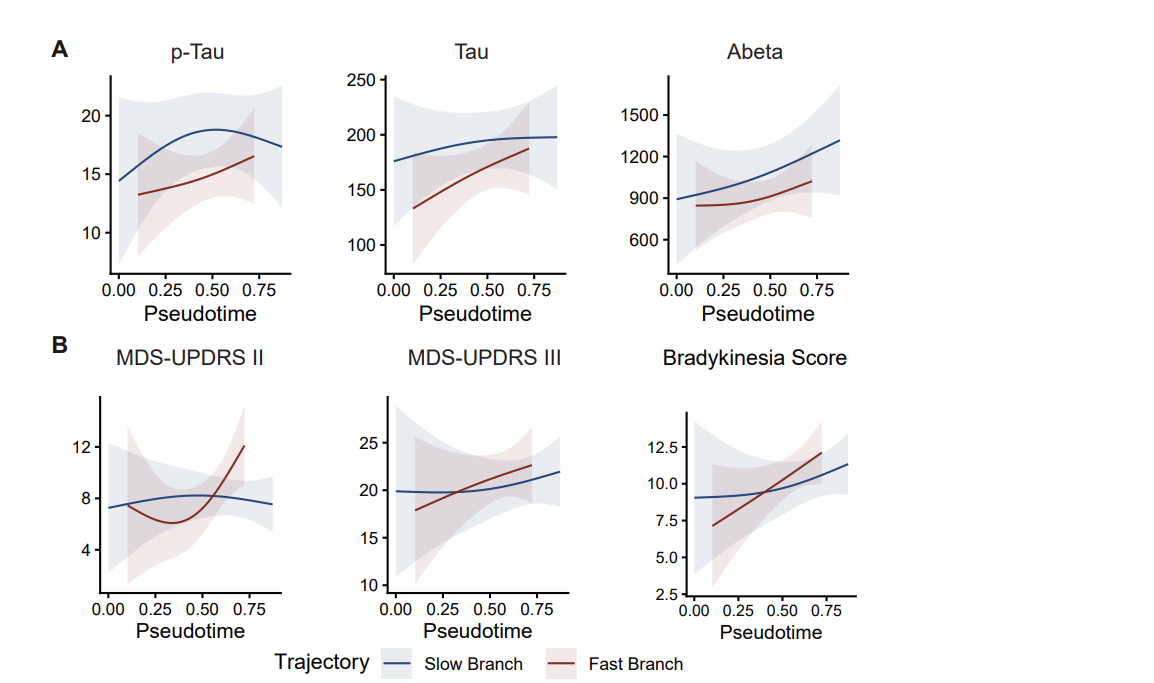

**Supplementary Figure 7. Application of the MOUSEPAD framework to female PD patients.** (**A**) Longitudinal trajectories of motor and non-motor clinical measures in female patients stratified by the MOUSEPAD framework. Female patients assigned to the fast-progression branch exhibited more rapid deterioration of clinical phenotypes compared with those assigned to the slow-progression branch. (**B**) Longitudinal trajectories of cerebrospinal fluid (CSF) biomarkers, including amyloid-β (Aβ), total Tau (t-Tau), and phosphorylated Tau (p-Tau), in female patients assigned to the fast- and slow-progression branches.

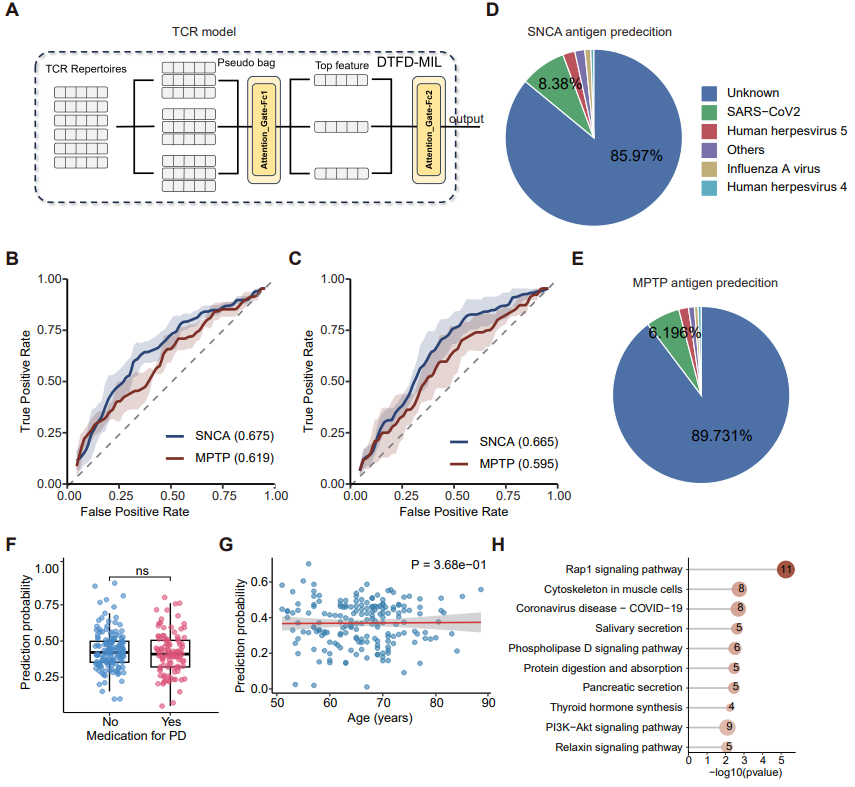

**Supplementary Figure 8. Deep learning–based TCR classification and biological interpretation.** (**A**) Schematic diagram illustrating the architecture of the deep learning model used for TCR-based classification. (**B-C**) Independent validation of model performance enhancement using diverse negative controls. These panels show the classification performance (ROC curves and AUCs) when training the model with additional samples from B) a COPD cohort or C) a CMV cohort as negative controls. Both additions independently improved performance, supporting the combined model presented in Fig. 5D. (**D-E**) Putative antigen prediction for key TCR features using TCR-match. Pie charts display the predicted antigen specificities for the top TCR features identified in the D) SNCA branch and E) MPTP branch. (**F**) Correlation between model prediction scores and patient medication status in the MPTP branch. (**G**) Correlation between patient age and model prediction score. (**H**) KEGG pathway enrichment analysis for the top 200 RNA features of the MPTP branch.
